# Evolution of Reduced Co-Activator Dependence Led to Target Expansion of a Starvation Response Pathway

**DOI:** 10.1101/101154

**Authors:** Bin Z. He, Xu Zhou, Erin K. O’Shea

## Abstract

In *S. cerevisiae*, the phosphate starvation (PHO) responsive transcription factors Pho4 and Pho2 are jointly required for induction of phosphate response genes and survival in phosphate starvation conditions. In the related human commensal and pathogen *C. glabrata*, Pho4 is required but Pho2 is dispensable for survival in phosphate-limited conditions and is only partially required for inducing the phosphate response genes. This reduced dependence on Pho2 evolved in *C. glabrata* and closely related species. Pho4 orthologs that are less dependent on Pho2 induce more genes when introduced into the *S. cerevisiae* background, and Pho4 in *C. glabrata* both binds to more sites and induces more genes with expanded functional roles compared to Pho4 in *S. cerevisiae*. Our work reveals an evolutionary mechanism for rapidly expanding the targets of a transcription factor by changing its dependence on a co-activator, potentially refining the physiological response it regulates.

## Introduction

For single celled organisms such as yeast, survival depends on stress response and signaling pathways that monitor the environment and generate appropriate responses to maintain homeostasis. How stress response pathways differ between species, especially in cases where the species occupy different environments, is of particular interest, because it illustrates the strategies used by an organism to adapt to its environment, and also reveals the evolutionary properties of a gene regulatory network as it diverges from a common ancestor. To approach this topic, we investigated the evolution of the phosphate starvation response (PHO) pathway between the model yeast S. *cerevisiae* and a related human commensal and opportunistic pathogen, *C. glabrata.* These two species are well suited for comparative study of stress responses given that they occupy distinct niches but still share ˜90% of their gene repertoire and have an average of ˜60% protein sequence identity (Gabaldón et al. 2013).

Phosphate is an essential nutrient for all organisms. To maintain phosphate homeostasis, S. *cerevisiae* activates a phosphate starvation pathway in response to limitation for inorganic phosphate (Ogawa, DeRisi, and Brown 2000). In phosphate-replete conditions, the transcription factor Pho4 is localized to the cytoplasm and phosphate response genes (PHO genes) are not expressed (O’Neill et al. 1996). As the concentration of extracellular inorganic phosphate (Pi) drops, cells activate the phosphate starvation response and Pho4 is imported into the nucleus where it functions together with the homeodomain transcription factor Pho2 to activate PHO gene expression (O’Neill et al. 1996; Komeili and O’Shea 1999; Vogel, Hörz, and Hinnen 1989; Barbaric et al. 1996; Barbaric et al. 1998; Shao, Creasy, and Bergman 1996).

Although Pho4 binds to ˜100 locations in the S. *cerevisiae* genome, it regulates fewer than 30 genes (Zhou and O’Shea 2011). Only genes at which Pho2 and Pho4 bind cooperatively in the promoter region are activated, indicating that Pho2 increases the selectivity of the gene set induced in response to phosphate starvation (Zhou and O’Shea 2011). In *C. glabrata*, Pho4 and Pho2 orthologs (hereinafter referred to as CgPho4 and CgPho2) exist, but unlike Pho4 and Pho2 in S. *cerevisiae* (hereinafter referred to as ScPho4 and ScPho2), CgPho4 can induce gene expression in the absence of CgPho2 (Kerwin and Wykoff 2009). This change in the dependence on the co-activator is not due to a higher expression level of CgPho4 or changes in the promoter regions of its target genes, and therefore is likely the result of alterations in the function of CgPho4 (Kerwin and Wykoff 2009).

We investigated the evolution of the PHO pathway in a diverse group of yeast species known as Hemiascomycetes (Knop 2006; Diezmann et al. 2004), which includes but is not limited to S. *cerevisiae, C. glabrata, K. lactis, C. albicans* and *Y. lypolitica*, and found that *PHO4* and *PHO2* are conserved as single copy genes in this group. We first evaluated Pho4 orthologs from a representative set of this group of species for their ability to activate gene expression in the absence of Pho2 in the S. *cerevisiae* background – this allowed us to establish that the reduced dependence on Pho2 evolved in a species clade that includes *C. glabrata*, two other human commensal yeasts and an environmental species. We then used functional genomics to assess the consequence of reduced Pho2 dependence on gene expression. Finally, we identified the *bona fide* targets of Pho4 in *C. glabrata* and compared them to Pho4 targets in S. *cerevisiae.* Our results show that gene regulatory networks can undergo target expansion by reducing the dependence of the major transcription factor on the co-activator, and as a result may lead to a new physiological response to the stress.

## Results

### Evolution of Pho2-dependence among Pho4 orthologs in the Hemiascomycetes

We first confirmed the previously reported result that deleting *PHO2* in *C. glabrata* does not eliminate expression of the secreted phosphatase encoded by *PMU2* ((Orkwis et al. 2010), Figure 1A), and then demonstrated that, in contrast to ScPho2, CgPho2 is dispensable for survival in phosphate-limited conditions (Figure 1B). To understand whether dependence on Pho2 in S. *cerevisiae* is the ancestral or the derived state and how this property of Pho4 evolved among related species, we surveyed the activity of Pho4 orthologs from 16 species in the Hemiascomycete class. To isolate the changes in Pho4 activity from the genomic background (e.g. promoter and Pho2 changes), we inserted coding sequences (CDSs) of the 16 Pho4 orthologs into an S. *cerevisiae* background lacking both the endogenous ScPho4 CDS and the negative regulator of the PHO pathway, Pho80.

**Figure 1.**
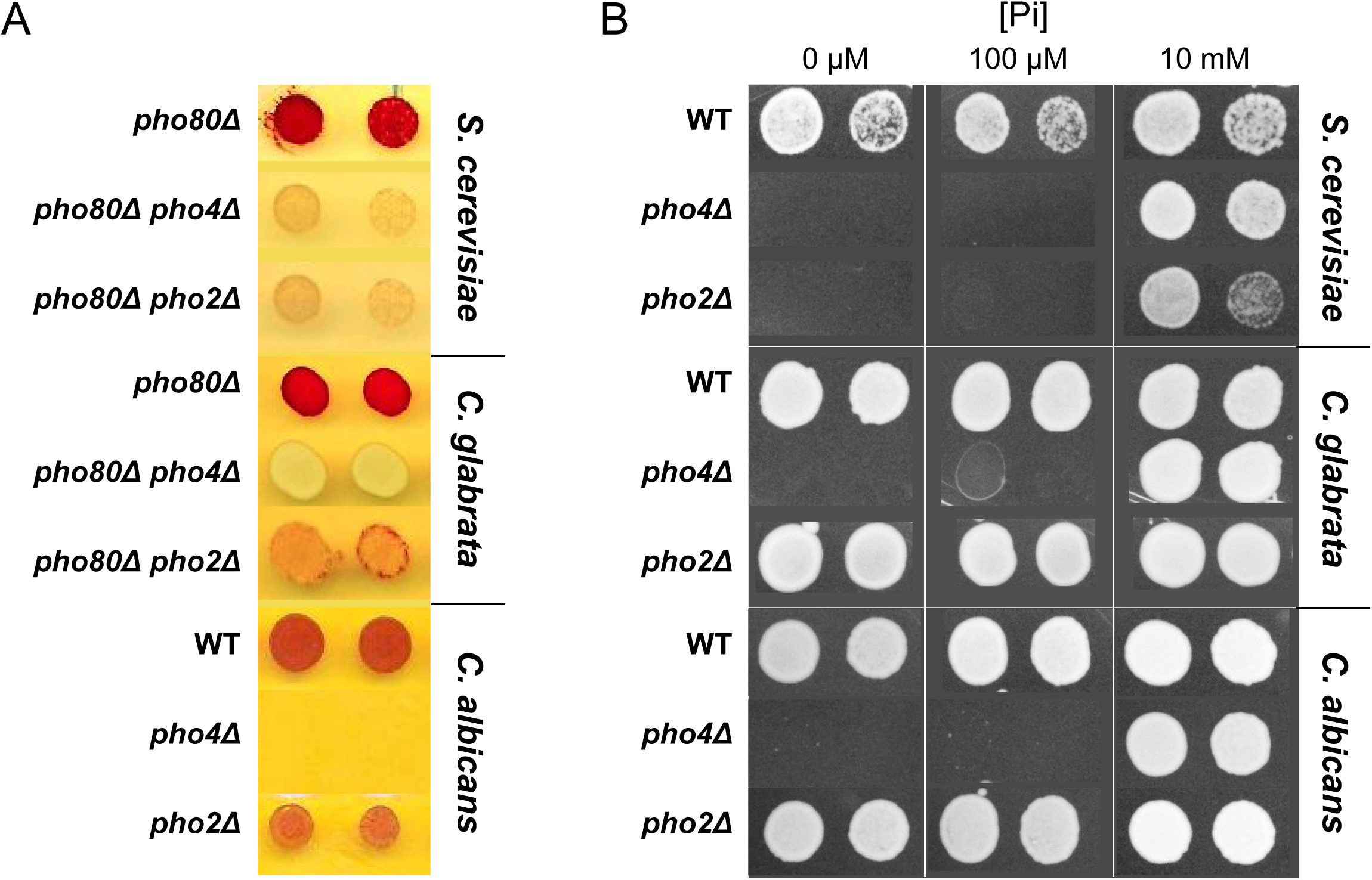
Difference among three yeast species in their dependence on Pho2 for gene induction and organismal survival under low Pi conditions. (A) Induction of the secreted phosphatase in each species measured by a semi-quantitative acid phosphatase assay (Wykoff 3t al. 2007). The intensity of the red color indicates the total activity of the secreted acid phosphatase from the cell colony. For S. *cerevisiae* and *C. glabrata*, strains lacking the negative regulator of Pho4 – Pho80 – were spotted on synthetic medium with 10 mM Pi. For *C. albicans*, strains with *PHO80* wild-type were spotted on synthetic medium lacking inorganic phosphate; the *pho2*Δ strain was not able to grow on this plate. (B) Colony growth phenotype of the wildtype, *pho4A*, *pho2*Δ strains in each of the three species, under different Pi concentrations. In both panels, two technical replicates of four-fold serial dilutions from the same culture are shown for each strain.

To determine the level of dependence on Pho2 for each Pho4 ortholog, we compared its activity, reflected by induction of the secreted phosphatase encoded by *PHO5*, in the presence or absence of ScPho2. We first evaluated whether the Pho4 orthologs can functionally compensate for ScPho4 in the S. *cerevisiae* background lacking the Pho4 negative regulator Pho80 and paired with ScPho2 (Figure 2A, left 3 columns). We found that the majority of Pho4 orthologs were able to induce Pho5 to a level significantly above the background *(pho4Á*, bottom row in Figure 2A), although the activity declines with increasing evolutionary distance from S. *cerevisiae.* When we measured the activity of the Pho4 orthologs in the absence of Pho2 (Figure 2A, right three columns), we found that Pho4 orthologs from the clade consisting of *C. glabrata, C. bracarensis, N. delphensis* and *C. nivariensis* (hereafter the *"glabrata* clade") were able to induce Pho5 expression in the absence of Pho2 (Figure 2A, B), but Pho4 orthologs from outgroup species such as *N. castellii, K. lactis*, and *L. waltii* could not. We conclude that the common ancestor of S. *cerevisiae* and *C. glabrata* had a Pho4 that is dependent on Pho2 and that the reduced dependence was evolutionarily derived in the *glabrata* clade.

**Figure 2.**
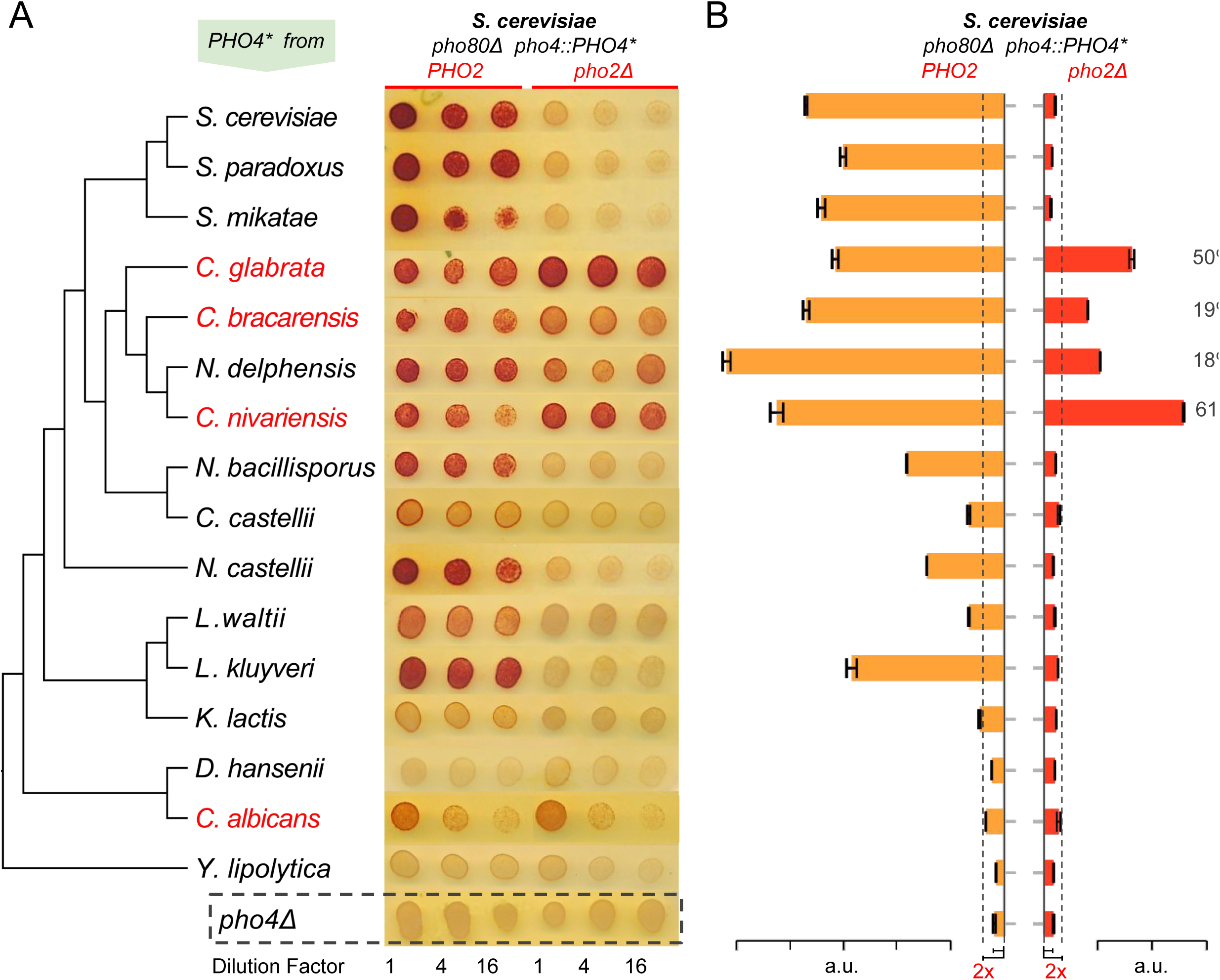
Evolution of Pho4 dependence on Pho2 in the Hemiascomycetes. (A) Survey of Pho4 orthologs activity in the S. *cerevisiae* background by the semi-quantitative acid phosphatase assay with or without ScPho2. The species phylogenetic relationship shown on the left were based on (Wapinski et al. 2007). Species names marked in red indicate known commensal and human pathogens. All strains were constructed in an identical S. *cerevisiae* background lacking the PHO pathway negative regulator Pho80. For each strain, three technical replicates in four-fold serial dilutions were assayed. A strain lacking Pho4 serves as the negative control (*pho4Δ*, dotted box). (B) Quantitative phosphatase assay for the same strains in (A). The bar graph shows the mean and standard deviation of the secreted phosphatase activity (N=2, technical replicates). For Pho4 orthologs with noticeable activities (exceeding twice the value of the *pho4Á* control with and without ScPho2, dotted lines), a percentage value was calculated by dividing the activity without ScPho2 by that with ScPho2, after subtracting the negative control (*pho4Δ*) in both cases. All results are representative of multiple (> 2) experiments.

Surprisingly, the Pho4 ortholog from a distantly related commensal and pathogenic yeast *C. albicans* (hereinafter “Ca-) weakly induced Pho5 in a Pho2-independent manner (Figure 2A).

We further demonstrated that *C. albicans* does not require Pho2 for survival in phosphate-limited conditions, and deletion of Pho2 does not abolish the induction of the secreted phosphatase in that species (Figure 1). We were not able to infer whether the reduced Pho2 dependence in *C. albicans* represents the derived or the ancestral state, because the Pho4 ortholog from *Y. lipolytica*, an outgroup of *C. albicans* and S. *cerevisiae*, failed to complement ScPho4 in S. *cerevisiae.*

In total, we identified five Pho4 orthologs with reduced dependence on Pho2, four of which come from known human commensal and opportunistic pathogens. This coincidence, while intriguing, is not sufficient to establish a true correlation between changes in PHO regulation and a commensal lifestyle, because the phylogenetic pattern represents at most two independent evolutionary events towards reduced Pho2 dependence, which could happen by chance alone. A broad phylogenetic sampling coupled with functional studies is required to establish or disprove this correlation.

### Reduced dependence on Pho2 is correlated with an increase in the number of Pho4 activated genes in *S. cerevisiae*

Since dependence on Pho2 provides additional selectivity for Pho4 induced gene expression in S. *cerevisiae* (Zhou and O’Shea 2011), we hypothesized that a reduction in Pho2-dependence will result in an increase in the number of Pho4 targets in the S. *cerevisiae* background. To test this prediction, we quantified the number of genes induced by different Pho4 orthologs expressed in an S. *cerevisiae* background lacking the negative regulator Pho80. We identified a total of 247 genes that were significantly induced by at least one of the eight Pho4 orthologs in the presence of ScPho2 (False discovery rate < 0.05, fold change > 2). Pho4 from *C. glabrata, C. bracarensis, N. delphensis* and *C. nivariensis* induced more genes than the Pho2-dependent Pho4 orthologs did (Figure 3A, D), and genes induced by these Pho4 orthologs are largely unaffected when ScPho2 is absent (Figure 3B). For example, 212 genes were induced by CgPho4 with ScPho2, compared to 40 genes induced by ScPho4 in the same background. Pho4 from S. *paradoxus*, a close relative of S. *cerevisiae*, induced a smaller number of genes than ScPho4, as did Pho4 from *L. kluyveri*, an outgroup of both S. *cerevisiae* and *C. glabrata.* Thus, the observed target expansion is not congruent with the phylogenetic relationship, but is a property unique to Pho4 orthologs with reduced Pho2-dependence. Moreover, differences in the mean expression levels of the Pho4 orthologs are small (< 2.5 fold, Figure 3- figure supplement 1) and do not explain the variation in their activity or dependence on Pho2 (Figure 3-figure supplement 2).

**Figure 3.**
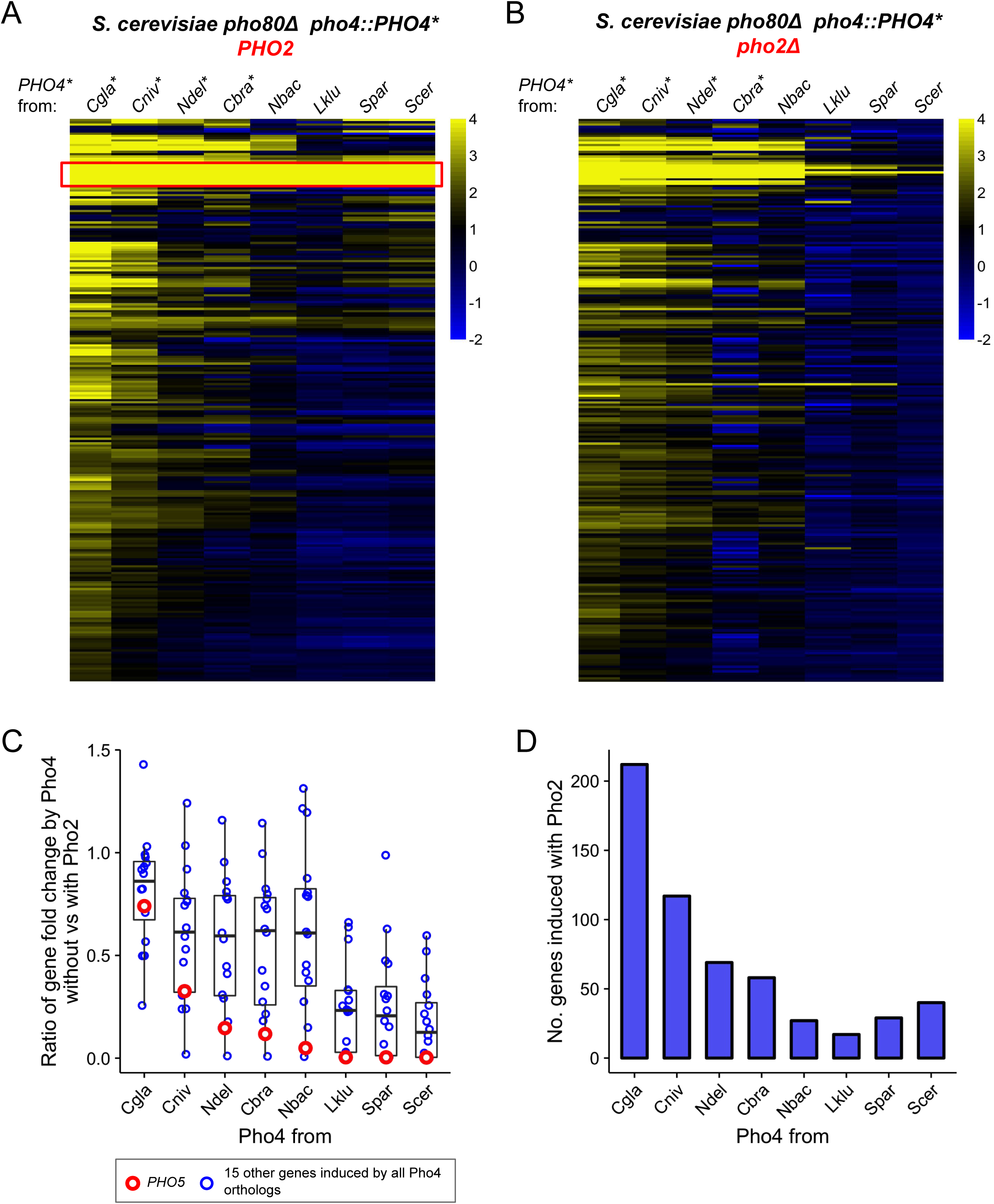
Pho4 orthologs that are less dependent on Pho2 induce more genes in the *S. cerevisiae* background. (A) Heatmap showing log_2_ fold change of genes (rows) induced by Pho4 orthologs (columns) in the S. *cerevisiae* background lacking *PHO80*, with ScPho2. A cutoff of 4 and −2 are used for visual presentation. The raw fold change estimates for the 247 genes by eight Pho4 orthologs were available in Figure 3- source data 1. Species names for each of the Pho4 orthologs were abbreviated and correspond to the full names in Figure 2. An asterisk indicates the Pho4 ortholog was shown to induce Pho5 expression in the absence of ScPho2 in S. *cerevisiae.* A total of 247 genes are plotted. The red box highlights a group of 16 genes that were induced by all eight Pho4 orthologs tested. (B) Same as (A) except the strains were in a *pho80Δ pho2*Δ background for all Pho4 orthologs. (C) Scatter plot comparing the levels of Pho2-dependence for each Pho4 ortholog, measured by the ratios for the 16 shared target genes between their fold changes in the absence versus in the presence of ScPho2. The boxplots represent the interquartile range (IQC, box), the mean (thick bar in the middle) and the highest or lowest values within 1.5 times of IQC (whisker). The red circles highlight one of the 16 genes, *PHO5.* (D) Bar plot showing the number of genes significantly induced more than twofold by each Pho4 ortholog in the presence of ScPho2.

We further investigated whether there is a quantitative relationship between the level of Pho2- dependence and the number of genes induced by Pho4 orthologs. We previously measured the level of Pho2-dependence by comparing the activity of a single Pho4 target – *PHO5* – in a pair of strains differing in the presence or absence of ScPho2. Here we made the same comparison for gene induction fold changes in a group of 16 genes induced by all eight Pho4 orthologs (Figure 3A, red box). The results, measured by the mean of the ratios for the 16 genes, are largely consistent with what we observed with Pho5 alone (Figure 3C), and, in general, the number of genes induced by each Pho4 ortholog increases with decreasing levels of Pho2- dependence (Figure 3D). In summary, the level of Pho2-dependence is negatively correlated with the number of genes induced by the Pho4 ortholog in the S. *cerevisiae* background.

### CgPho4 binds to more locations in *S. cerevisiae* and activates a higher percentage of genes upon binding

We reasoned that the expansion of target genes for the *glabrata* clade Pho4 orthologs could result from Pho4 binding to more genomic locations, Pho4 activating a higher proportion of the genes to which it binds, or a combination of the two. To test if differences in Pho4 binding account for target gene expansion, we performed chromatin immunoprecipitation followed by high-throughput sequencing (ChIP-seq) to identify the binding locations for both ScPho4 and CgPho4 in the S. *cerevisiae* background lacking the negative regulator Pho80. We identified a total of 115 ChIP-peaks for CgPho4 and 74 peaks for ScPho4, with 72 being bound by both (Figure 4A, Figure 4- source data 1). The expansion of CgPho4 binding locations was not because it recognized new sequence motifs – 42 of the 43 CgPho4-specific peaks contain the consensus “CACGTG- motif, and 69 of the 72 shared peaks contain this motif. For all four exceptions, a one-base-pair (bp) mismatch to the consensus motif is observed (Figure 4A, parentheses). Therefore, DNA binding specificity is conserved between ScPho4 and CgPho4. In contrast, ScPho4 and CgPho4 differ in their dependence of DNA binding on Pho2 – ScPho4 binding is significantly lower when ScPho2 is absent, but CgPho4 binding is largely unaffected by the deletion of ScPho2 (Figure 4B). In summary, CgPho4 recognizes the same E-box motif as ScPho4 does, but its binding is no longer dependent on ScPho2 and CgPho4 binds to ˜50% more (43/74 ≈ 0.55) sites than ScPho4 does.

**Figure 4.**
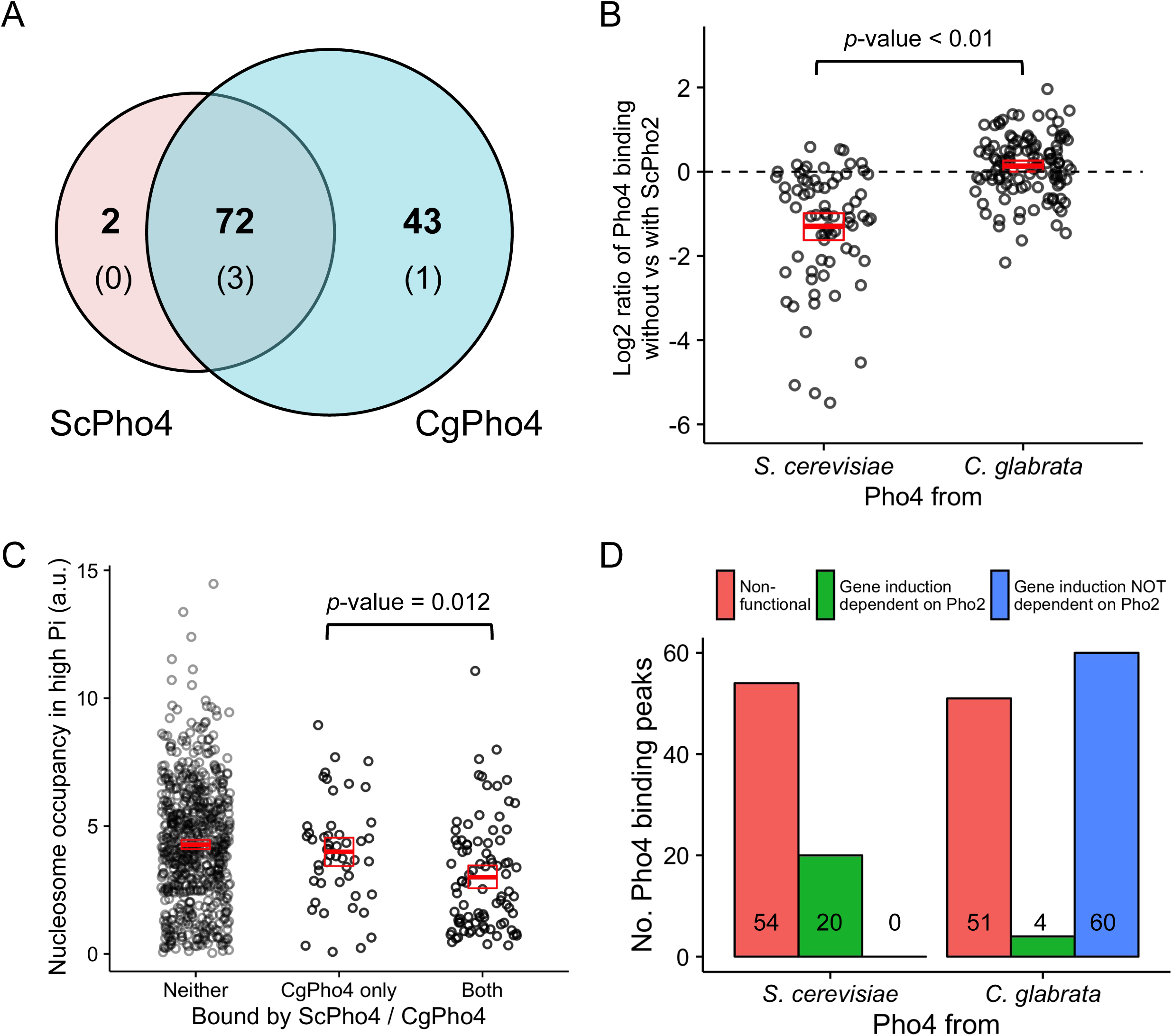
CgPho4 binds to more genomic locations than ScPho4 and is more likely to lead to gene activation upon binding in *S. cerevisiae*. (A) Venn diagram showing the number of and overlap between binding locations for ScPho4 and CgPho4 in the S. *cerevisiae* genome. The numbers in parentheses indicate binding events among the total number where the DNA sequence underlying the peak contains a suboptimal motif with one base pair mismatch to the consensus. (B) Scatter plot showing log_2_ ratio of ScPho4 or CgPho4 ChIP occupancy without vs with ScPho2. Only sites bound by Sc or CgPho4 in the presence of ScPho2 (N = 74 and 115, respectively) are plotted. The thick red bar represents the mean and the box the 95% confidence limits computed by a non-parametric bootstrapping method (Harrell 2016). The means of the two groups are significantly different by a two-sided Student’s t-test, with a p-value < 0.01. (C) Scatterplot for nucleosome occupancy in high Pi conditions at “CACGTG- motifs either bound by neither ScPho4 nor CgPho4 (N = 660), only by CgPho4 (N = 48) or by both (N = 88). The red bar and box have the same meaning as in (B), and the difference between sites bound only by CgPho4 and those bound by both CgPho4 and ScPho4 is significant by a two-sided Student’s t test (<u>*p*</u>-value = 0.012). (D) Bar plot comparing the number of genome-wide binding peaks for ScPho4 and CgPho4 that are either non-functional, lead to gene induction only with ScPho2 or lead to gene induction with or without ScPho2. The source data listing all identified ChIP peaks and the associated gene induction statistics are provided in Figure 4- source data 1.

It has been reported that both nucleosomes and another transcription factor with similar binding specificity to Pho4, Cbf1, competitively exclude ScPho4 from the E-box motifs in the genome (Zhou and O’Shea 2011). It is therefore plausible that CgPho4 is able to bind to more locations because it can access sites normally occupied by nucleosomes or competitors. To test if nucleosome exclusion plays a role, we mapped the published nucleosome occupancy in phosphate-replete (high Pi) conditions (Zhou and O’Shea 2011), where Pho4 is inactive, to the binding peaks identified in this study, to find that sites bound only by CgPho4 have on average higher nucleosome occupancy in high Pi conditions than sites bound by both CgPho4 and ScPho4 (Figure 4C, t-test for difference in the mean: p-value = 0.012, two-sided test). We performed the same analysis for Cbf1 enrichment in high Pi conditions at the top 25% most accessible E-box motifs – those with the lowest nucleosome occupancy – to avoid confounding nucleosome competition with Cbf1 binding. We found that Cbf1 enrichment at sites bound only by CgPho4 is not significantly different from enrichment at sites bound by both ScPho4 and CgPho4 (Figure 4- figure supplement 1). It is worth noting, however, that the small sample size in the CgPho4 bound only class (11) may have precluded us from detecting small differences. In conclusion, CgPho4 binding is less dependent on ScPho2 and it competes more favorably with nucleosomes than ScPho4 does, which likely contributes to the expansion of CgPho4 binding sites in the S. *cerevisiae* genome.

To compare the ability of CgPho4 and ScPho4 to induce gene expression upon binding to the promoter, we analyzed the transcriptional profiling data for the genes bound by the two Pho4 orthologs. We found that CgPho4 not only bound to more sites, but it also activated a higher percentage of the downstream genes upon its binding than did ScPho4 (64/115 = 56% vs 20/74 = 29%, Figure 4D). Moreover, its ability to induce gene expression is largely independent of Pho2: >90% of CgPho4 targets (60/64 = 93.75%) were induced in the *pho2Δ* background, while all (20/20) ScPho4 induced genes required ScPho2 (Figure 4D).

In conclusion, CgPho4 both binds DNA and activates gene expression independently of ScPho2. Compared to ScPho4, it is more capable of accessing nucleosome-occluded binding motifs and it is also able to activate downstream gene expression with a higher probability upon binding. We propose that the combination of these features led to the expansion in the targets of CgPho4 in S. *cerevisiae* and speculate that this may underlie the target expansion for Pho4 from *C. bracarensis, N. delphensis* and *C. nivariensis.*

### In *C. glabrata* CgPho4 binds to DNA and activates gene expression largely independent of Pho2

Next we asked if CgPho4 also functions independently of Pho2 in its endogenous genome. To investigate the dependence of CgPho4 binding on CgPho2, we used the high-resolution ChlP-exo technique to map CgPho4 binding in the presence and absence of CgPho2, and CgPho2 binding, under both phosphate-replete and phosphate-limited conditions (Materials and Methods) (Rhee and Pugh 2012. We identified a total of 100 binding peaks for CgPho4 in the presence of CgPho2 under phosphate-limited conditions (Figure 5- source data 1). CgPho4 recognizes the same “CACGTG- motif as it does in S. *cerevisiae* (Materials and Methods), and the consensus motif is present in 51 of the 100 peaks, with the rest containing a one-bp mismatch (46) or two-bp mismatches (3). CgPho2 bound to more than 500 sites genome-wide, without a strongly enriched sequence motif (Materials and Methods). With respect to CgPho4 bound sites, CgPho2 binds at the same location for 77 of the 100 peaks (Figure 5A). Among these shared binding peaks, only 14 (18%) CgPho4 peaks showed more than two-fold reduction in peak height in the *pho2*Δ background (Figure 5A). We hypothesized that the quality of the DNA motif underlying the peak may explain the differential requirement of Pho2 co-binding. We tested this hypothesis by comparing changes in CgPho4 binding when CgPho2 is deleted, at sites with a consensus motif vs those without (Figure 5- figure supplement 1). Although the trend matches our expectation, the difference is small and not significant at a 0.05 level by the Student’s t-test (p-value = 0.11). We conclude that CgPho4 binding to DNA is largely independent of CgPho2 in *C. glabrata*, but a small fraction (18%) of its binding sites show CgPho2 influence.

**Figure 5.**
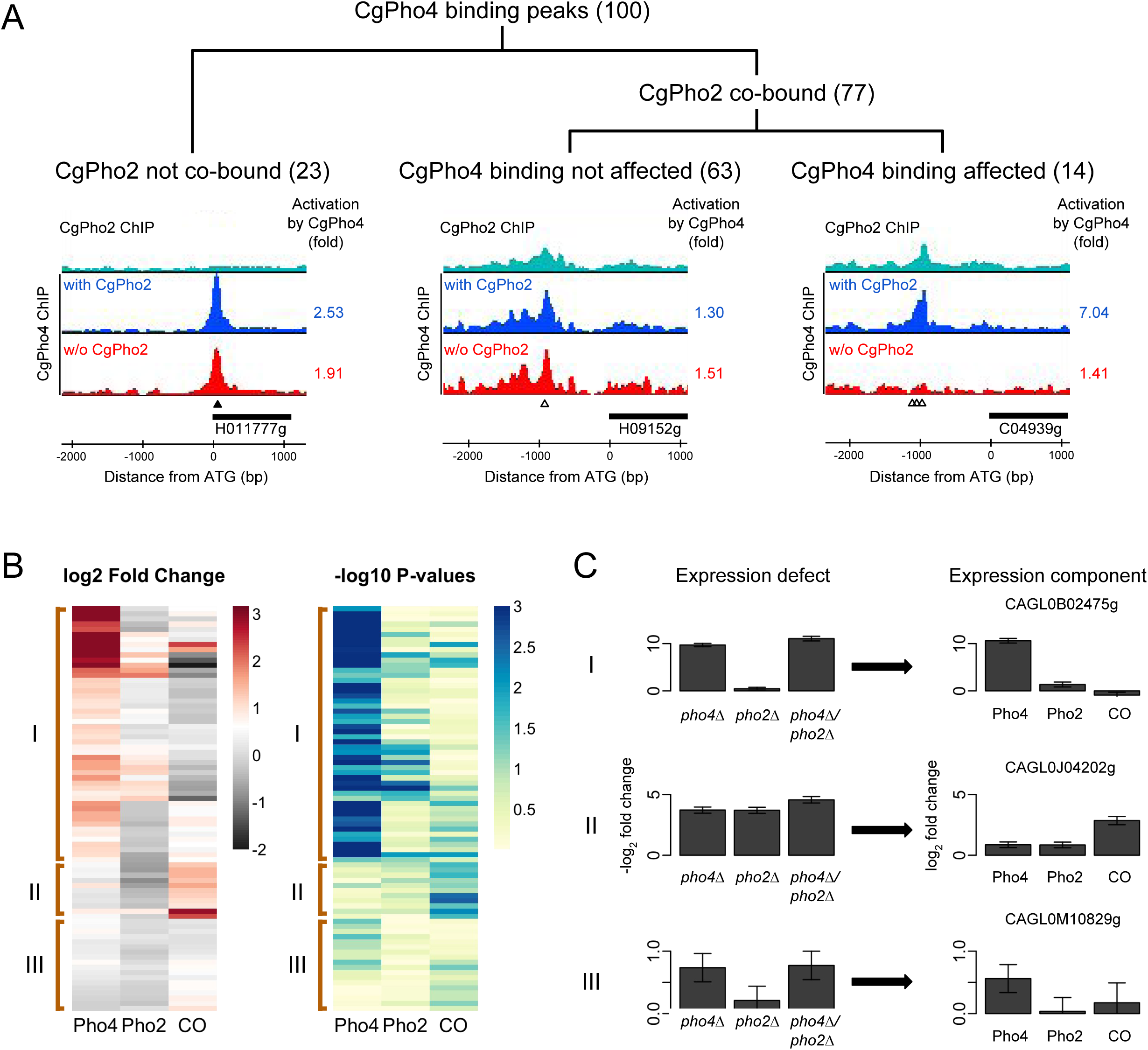
Identifying Pho4 targets in *C. glabrata* using genome-wide binding and transcriptome profiling data. (A) Dendrogram showing the breakdown of the 100 CgPho4 ChIP peaks based on whether CgPho2 binds next to CgPho4, and when it does, whether CgPho4 binding is affected by the deletion of Pho2 or not (defined as CgPho4 ChIP peak height reduced by more than twofold in the *pho2*Δ background). The graphs below the dendrogram show examples of ChIP profiles for each category of CgPho4 binding. Profiles of ChIP fold enrichment over mock are shown for CgPho2 in cyan, CgPho4 with CgPho2 in blue and CgPho4 without CgPho2 in red. The downstream gene is depicted as a thick bar to the right and the shortened systematic name (remove the preceding “CAGL0-) is shown below. The fold changes in induction for the putative target gene with or without CgPho2 are shown to the right of each graph. A list of the 100 binding peaks and the associated statistics are available in Figure 5- source data 1. (B) The left heat map showing the estimates of expression components for 79 genes directly bound and induced by CgPho4. For each gene, the log_2_ transformed fold change is decomposed into Pho4 effect alone (Pho4) + Pho2 effect alone (Pho2) + Pho4/Pho2 collaborative effect (CO). A cutoff of 3 and −2 is used for visual presentation. The unadjusted estimates and the associated p-values are available in Figure 5- source data 2. The right heatmap shows the −log_10_ transformed p-values for the corresponding t-statistics of the estimates on the left. Three groups are defined based on their characteristic expression components: group I genes are dominated by CgPho4 main effect; group II genes depend on both CgPho4 and CgPho2 (CO component); group III genes are a mix of the first two groups, with lower fold changes (weakly induced). (C) Bar graphs on the left showing the linear model estimates and the standard deviation of the expression defects, defined as the −log2 transformed fold changes between the mutants (single or double) and the wild-type. On the right are the corresponding estimates and standard deviation of the expression components for the same gene, estimated from the same data with two biological replicates per strain (Materials and Methods). One representative gene is plotted for each category in (B).

To evaluate whether gene induction by CgPho4 is dependent on CgPho2, we used RNA-seq to quantify the fold changes for genes induced by CgPho4 with or without CgPho2, in a strain lacking the negative regulator CgPho80. Intersecting with the ChIP identified CgPho4 binding sites, we identified 79 genes that were both directly bound and induced by CgPho4 in the presence of CgPho2 (Figure 5-source data 2). We then used mutant-cycle analysis (Capaldi et al. 2008) to delineate the contribution from either CgPho4 acting alone (Pho4), CgPho2 acting alone (Pho2) or the two factors acting cooperatively (CO), and used unsupervised clustering (Ward’s method, Materials and Methods) on the estimated values for the three components to group the genes into three classes (Figure 5B,C). In class I (50 genes), CgPho4 is the dominant contributor to gene induction, with CgPho2 either contributing to a lesser extent by itself (Pho2) or through its interaction with CgPho4 (CO). Class II (11) genes show a strong collaborative component, with little contribution from Pho4 acting alone. Class III (18) genes show relatively low fold changes, with the main contribution coming from either CgPho4 acting alone or its collaborative effect with CgPho2. In conclusion, we found that >60% (50/79) of the genes bound by CgPho4 are induced primarily by CgPho4 acting alone, and that a lesser fraction (11/79) depend on the collaborative action of CgPho4 and CgPho2. This is in contrast to S. *cerevisiae*, where the majority of gene induction was attributed to the cooperative interaction between ScPho4 and ScPho2 (23/28 genes are induced only when both ScPho4 and ScPho2 are present (Zhou and O’Shea 2011)).

### #Pho4 direct targets in *C. glabrata* may function beyond phosphate homeostasis

In S. *cerevisiae*, nearly all ScPho4 targets function in either regulating the PHO pathway or maintaining intracellular phosphate homeostasis (Ogawa, DeRisi, and Brown 2000; Zhou and O’Shea 2011). To gain insight into the function of the PHO pathway in *C. glabrata*, we studied the Gene Ontology (GO) terms associated with the 79 genes bound and induced by CgPho4 in *C. glabrata* (Figure 5- source data 2). The top three enriched GO terms for CgPho4 targets are related to phosphate homeostasis, i.e. polyphosphate metabolism, phosphorus metabolism and phosphate ion transport (Figure 6- source data 1), confirming that the PHO pathway in *C. glabrata* is conserved in its core function. However, the genes in this core functional group are not all conserved (Figure 6A). Underlying the apparent conservation in function are non-orthologous genes that are either paralogs (e.g. *HOR2* in S. *cerevisiae* vs *RHR2* in *C. glabrata)* or evolutionarily unrelated (e.g. the phosphatase function of *PHO5* in S. *cerevisiae* is replaced by that of *PMU2* in *C. glabrata)* (Figure 6A, Figure 6- source data 2, (Kerwin and Wykoff 2009; Kerwin and Wykoff 2012; Orlando et al. 2015)).

**Figure 6.**
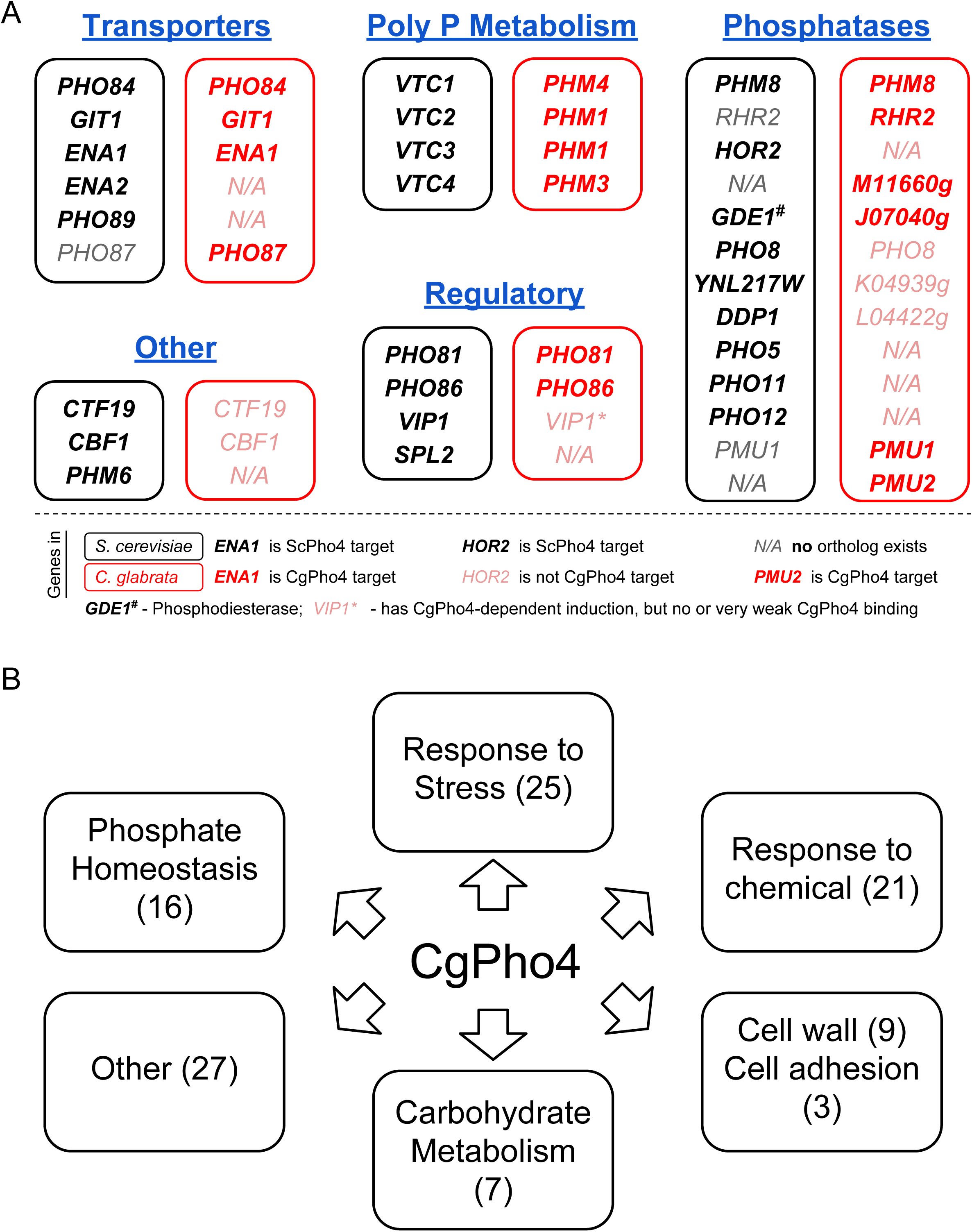
Functional annotation of Pho4 targets in *C. glabrata*. (A) Comparison between all 24 ScPho4 targets in S. *cerevisiae* and CgPho4 targets with phosphate homeostasis related functions in *C. glabrata.* Abbreviations: “Poly P- stands for “Polyphosphate-. Within each subcategory, S. *cerevisiae* genes in black are paired with their homologs in *C. glabrata* in red. N/A in either species indicates ortholog does not exist in that species. *C. glabrata* genes not annotated with a common name are represented by their systematic name with the preceding “CAGL0- omitted. Gene names in bold indicate that they are targets of ScPho4 or CgPho4, while gray or pink gene names are not targets of ScPho4 or CgPho4, respectively. (B) Non-exclusive groups of Pho4 targets in *C. glabrata* based on Gene Ontology “Biological process- terms and functional annotations in Candida Genome Database (for *C. glabrata* or orthologs in *C. albicans)* and Saccharomyces Genome Database (for orthologs in S. *cerevisiae).*

Genes with functions related to phosphate homeostasis account for only 16 of 79 CgPho4 targets. To predict the functions of the remaining CgPho4 targets, we mapped all 79 genes to GO Slim terms and identified several major functional groups (Figure 6B, Figure 6- source data 3). Pho4 targets in *C. glabrata* are enriched in genes predicted to be involved in the response to non-phosphate-related stresses, response to chemical stresses, fungal cell wall biosynthesis and cell adhesion, and carbohydrate metabolism (Figure 6- source data 4- Table S1-5). Thus, it appears that the PHO regulon in *C. glabrata* has expanded in the number of targets, and likely expanded its function beyond phosphate homeostasis. It is worth mentioning that, although CgPho4 induces more genes in S. *cerevisiae* as well as in *C. glabrata*, there is virtually no overlap between its targets in S. *cerevisiae* and in *C. glabrata*, except for those involved in phosphate homeostasis (as shown in Figure 6A). This is interesting because it may be intuitively appealing to assume that the additional targets of CgPho4 in *C. glabrata* were evolutionarily acquired by exploiting existing E-box motifs in the genome that were inaccessible to the ancestral Pho2-dependent Pho4. This may still be true for some of the targets, as it is possible that the S. *cerevisiae* orthologs of those *C. glabrata* genes lost the E-box motifs after the two species diverged. The alternative hypothesis is that the targets not involved in phosphate homeostasis were gained *de novo* in *C. glabrata* by their promoters acquiring CgPho4-recognized motifs. In conclusion, our comparative study of the Pho network between S. *cerevisiae* and *C. glabrata* reveals a substantial target expansion in the latter species. This comparison, unfortunately, does not provide the temporal resolution for reconstructing the tempo and mode of target expansion in *C. glabrata.* Comparative studies of more closely related species are needed to address this outstanding question.

## Discussion

### Evolutionary constraint and plasticity in the PHO response network

Phosphate is an essential macronutrient, and starvation for inorganic phosphate is likely a common stress experienced by all microorganisms. Hence it is not surprising that the PHO response is ancient and conserved among the Hemiascomycete yeasts. In particular, the transcription factors Pho4 and Pho2 are present in all 16 species with whole genome sequences, and Pho4 is known to regulate the PHO response in distantly related species such as *C. albicans* (Ikeh et al. 2016). Pho2 likely acts as the co-activator in the majority of the species according to our assay (Figure 2), and its role in assisting Pho4, although diminished, is still detectable in *C. glabrata* (Figure 5). Also conserved is the mechanism for regulating Pho4 activity in response to phosphate starvation – in both *C. glabrata* and S. *cerevisiae*, Pho4 nuclear localization is phosphorylation-dependent, controlled by homologous cyclin-dependent kinase complexes (Kerwin and Wykoff 2012). Finally, our ChIP results demonstrated that both ScPho4 and CgPho4 recognize the same E-box motif “CACGTG-.

In contrast to the transcription factor and its regulation, the downstream targets of the PHO response are much more evolutionarily dynamic. One mode of evolution, which we term “re-assignment-, involves non-orthologous genes performing the same or similar functions (Figure 6A). The gene counterparts are either paralogs, e.g. *ScGPP2* (=*HOR2*) vs *CgGPP1* (=*RHR2*), or they can be evolutionarily unrelated, e.g. the repressible acid phosphatase encoded by *ScPHO5*vs *CgPMU2*. In the latter case, the two acid phosphatases were shown to exhibit different substrate specificities – Pho5 has much higher activity for phytic acid, an organic phosphate-containing compound that is relatively abundant in plant materials (S. *cerevisiae* environment) but generally not available in mammalian tissues (C. *glabrata* environment) (Orkwis et al. 2010). Hence it is possible that this evolutionary reassignment may be driven by selection to optimize the PHO response for the specific environment the organism lives in.

Another mode of evolution involves expansion (or contraction (Ihmels et al. 2005)) of the gene targets. New targets can be incorporated into an existing network either via *de novo* gain of DNA motifs in the promoters (Gompel et al. 2005), or by changes in the transcription factor protein-DNA interaction (Cheatle Jarvela et al. 2014; McKeown et al. 2014). The former is less pleiotropic, albeit slow if a large number of new targets are to be incorporated (Wittkopp and Kalay 2011; Stern 2010). Evolution of the transcription factor protein, on the other hand, is highly constrained (Maerkl and Quake 2009; Struhl 1987; Nitta et al. 2015; Wilson et al. 2008), although changes in binding specificity and gain of secondary motifs have been observed (Sayou et al. 2014; Cheatle Jarvela et al. 2014; McKeown et al. 2014). We found that the DNA sequence preference is highly conserved between ScPho4 and CgPho4. Instead, evolution of reduced dependence on the co-activator allowed CgPho4 to rapidly expand its targets without affecting existing ones. Interestingly, the same molecular trait might have independently evolved in the distantly related commensal and pathogen, *C. albicans* (Figure 1, 2), and a recent study showed CaPho4 has an expanded target set (Ikeh et al. 2016). We postulate that changes in the dependence on the co-activator may be repeatedly used in evolution to rapidly expand the targets without affecting existing ones. More generally, changes affecting the interaction between transcription factors and their co-factors may be an important yet underappreciated mechanism underlying both gene regulatory network evolution and certain disease phenotypes (Slattery et al. 2011; Ang et al. 2016, 4). In conclusion, what we observed in the evolution of the PHO response network in *C. glabrata* is both conservative and flexible: it is conservative in the transcription factor identity and its regulation, but flexible in the downstream targets via changes in Pho4’s dependence on the co-activator Pho2.

### Expansion in the function of the PHO regulon and its implication for the stresses faced by *C. glabrata*

Only 16 of the 79 genes directly induced by CgPho4 in *C. glabrata* are involved in maintaining phosphate homeostasis. Among the remaining genes, a significant number (25 genes) are predicted to be involved in responses to non-phosphate related stresses, including osmotic (7 genes) and oxidative (7 genes) stresses. In addition to roles in other stress responses, a smaller group of CgPho4 targets (9 and 3 genes) have potential functions in cell wall synthesis and cell adhesion, two traits that were known to be relevant for survival and virulence in the host (Atanasova et al. 2013; De Las Peñas et al. 2015; Luo and Samaranayake 2002; Jawhara et al. 2012; Fabre et al. 2014). A similar observation has been made before in *C. glabrata*, where limitation of nicotinic acid was sufficient to induce genes mediating a cellular adhesion phenotype (Domergue et al. 2005). Intriguingly, Pho4 in *C. albicans* has been shown to be important for survival under particular types of osmotic and oxidative stresses, suggesting a similar functional expansion as we observed in *C. glabrata* (Ikeh et al. 2016). Also in *C. albicans*, phosphate starvation was linked to enhanced virulence, and a strain lacking Pho4 displayed extensive filamentation in response to phosphate limitation (Romanowski et al. 2012). Combining these observations, we speculate that stress responses in the commensal species may have evolved to be more coordinated, and may have acquired new targets linked to virulence, to cope with the distinct stress profiles in the host, such as spatiotemporally overlapping challenges exerted by the host immune cells (Kasper, Seider, and Hube 2015). Characterizing other stress response pathways in *C. glabrata*, and in other commensal species such as *C. albicans*, can test this hypothesis and will shed further light on the architectural differences in the stress and starvation response network compared to that in the free-living species.

## Methods

### Strains

#### S. cerevisiae strains

All S. *cerevisiae* strains were generated from EY0057 (K699 *ade2-1 trp1-1 can1-100 leu2-3,112 his3-11,15 ura3* GAL+), using a standard high efficiency transformation protocol with ˜40 bp homology sequences (Gietz and Schiestl 2007). The Pho4 ortholog swap strains (EY2863 etc., Table 1) were made by the *URA3* pop-in and pop-out method. Briefly, a *K. lactis URA3* gene with proper promoter and terminator was used to precisely replace the CDS of *PHO4* in EY2849, where *PHO80* is deleted. The *URA3* gene was then replaced by the CDS sequence from one of the *PHO4* orthologs, obtained via PCR-based cloning from the genomic DNA of the relevant species. Successful transformants were validated using PCR. *PHO2* was knocked out using *KlacURA3*, generating the complementary set of *pho2Δ* strains. Genomic DNA for non- *cerevisiae* species was prepared using standard methods (ball milling to break the cells, followed by Phenol Chloroform extraction and RNase treatment). PCR primers were designed based on genome sequences for each species from the Orthogroup website (Wapinski et al. 2007), except for *C. nivariensis, C. bracarensis, N. bacillisporus, N. delphensis* and *C. castellii*, which were initially provided to us by Dr. Cecile Fairhead and are now available through GRYC (Genome Resources for Yeast Chromosomes, http://gryc.inra.fr). Strains of the non-S. *cerevisiae* species were from Aviv Regev lab (Thompson et al. 2013) and Cecile Fairhead (for the *Nakaseomyces* genus). Abbreviations for species names are as follows: S. *mikatae (Smik), S. paradoxus (Spar), C. glabrata (Cgla), C. nivariensis (Cniv), C. bracarensis (Cbra), N. bacillisporus (Nbac), N. delphensis (Ndel), C. castellii (Ccas), N. castellii (Ncas), L. waltii (Lwal), L. kluyveri (Lklu), K. lactis (Klac), D. hansenii (Dhan), C. albicans (Calb), Y. lipolytica (Ylip).* Species names were based on YGOB and GRYC (Byrne and Wolfe 2005; Gabaldón et al. 2013). Bio-ChIP strains for either ScPho4 or CgPho4 were generated by replacing the *URA3* marker used to knockout the *PHO4* CDS with a linear DNA construct containing either ScPho4 or the CDS of CgPho4 fused with the Avitag (GLNDIFEAQKIEWHW) at the C-terminus, separated by a “GSGSGS- linker. Primer sequences for generating the strains are available upon request.

**Table 1.** Strains used in this Study

| Strain | Genotype | Reference |
| --- | --- | --- |
| ***S. cerevisiae*** |  |  |
| EY0057 | K699, as wild-type for PHO pathway | (O'Neill et al. 1996) |
| EY0252 | *pho2::LEU2* | (Kaffman et al. 1994) |
| EY1710 | *pho4::URA3* | (Zhou and O'Shea 2011) |
| EY2849 | *pho80::TRP1* | This study |
| EY2851 | *pho80::TRP1 pho2::HIS5* | This study |
| EY2852 | *pho80::TRP1 pho4::URA3* | This study |
| EY2859 | *pho80::TRP1 pho4::URA3 pho2::HIS5* | This study |
| EY2863 | *pho80::TRP1 pho4::CglaPHO4* | This study |
| EY2872 | *pho80::TRP1 pho4::SparPHO4* | This study |
| EY2873 | *pho80::TRP1 pho4::SmikPHO4* | This study |
| EY2874 | *pho80::TRP1 pho4::LkluPHO4* | This study |
| EY2875 | *pho80::TRP1 pho4::NcasPHO4* | This study |
| EY2876 | *pho80::TRP1 pho4::KlacPHO4* | This study |
| EY2877 | *pho80::TRP1 pho4::LwalPHO4* | This study |
| EY2878 | *pho80::TRP1 pho4::DhanPHO4* | This study |
| SC1 | *pho80::TRP1 pho4::CnivPHO4* | This study |
| SC2 | *pho80::TRP1 pho4::CbraPHO4* | This study |
| SC4 | *pho80::TRP1 pho4::NbacPHO4* | This study |
| SC5 | *pho80::TRP1 pho4::NdelPHO4* | This study |
| SC26 | *pho80::TRP1 pho4::CcasPHO4* | This study |
| SC33 | *pho80::TRP1 pho4::CalbPHO4* | This study |
| SC35 | *pho80::TRP1 pho4::YlipPHO4* | This study |
| EY2879 | *pho80::TRP1 pho4::CglaPHO4 pho2::URA3* | This study |
| EY2880 | *pho80::TRP1 pho4::SparPHO4 pho2::URA3* | This study |
| EY2881 | *pho80::TRP1 pho4::SmikPHO4 pho2::URA3* | This study |
| EY2882 | *pho80::TRP1 pho4::LkluPHO4 pho2::URA3* | This study |
| EY2883 | *pho80::TRP1 pho4::NcasPHO4 pho2::URA3* | This study |
| EY2884 | *pho80::TRP1 pho4::KlacPHO4 pho2::URA3* | This study |
| EY2885 | *pho80::TRP1 pho4::LwalPHO4 pho2::URA3* | This study |
| EY2886 | *pho80::TRP1 pho4::DhanPHO4 pho2::URA3* | This study |
| SC7 | *pho80::TRP1 pho4::CnivPHO4 pho2::URA3* | This study |
| SC8 | *pho80::TRP1 pho4::CbraPHO4 pho2::URA3* | This study |
| SC9 | *pho80::TRP1 pho4::NbacPHO4 pho2::URA3* | This study |
| SC10 | *pho80::TRP1 pho4::NdelPHO4 pho2::URA3* | This study |
| SC27 | *pho80::TRP1 pho4::CcasPHO4 pho2::URA3* | This study |
| SC34 | *pho80::TRP1 pho4::CalbPHO4 pho2::URA3* | This study |
| SC36 | *pho80::TRP1 pho4::YlipPHO4 pho2::URA3* | This study |
| EY2681 | *pho80::HIS3 pho4::ScerPHO4-C-AVI-TRP1 ura3::pRS306-BirA* | (Zhou and O'Shea 2011) |
| EY2869 | *pho80::TRP1 pho4::CglaPHO4-C-AVI ura3::pRS306-BirA* | This study |
| EY2867 | *pho80::TRP1 pho4::CglaPHO4-C-AVI pho2::URA3 ura3::pRS306-BirA* | This study |
| SC32 | *pho80::TRP1 pho4::CglaPHO4 ura3::pRS306-BirA* | This study |
| ***C. glabrata*** |  |  |
| BG99 | *his3Δ* (1 + 631), as wild-type for PHO pathway | (Cormack and Falkow 1999) |
| CG1 | BG99, *ura3Δ* | This study |
| CG2 | *pho80::HIS3* | This study |
| CG3 | *pho4::URA3* | This study |
| CG4 | *pho80::HIS3 pho4::URA3* | This study |
| CG5 | *pho2::URA3* | This study |
| CG6 | *pho80::HIS3 pho2::URA3* | This study |
| CG16 | *pho80::HIS3 pho4::natNT2 pho2::URA3* | This study |
| CG40 | *pho2::N-V5-PHO2* | This study |
| CG43 | *pho4::PHO4-C-3xFLAG* | This study |
| CG49 | *pho4::PHO4-C-3xFLAG pho2::URA3* | This study |
| CG51 | *pho80::HIS3 pho2::N-V5-PHO2* | This study |
| ***C. albicans*** |  |  |
| Wild-type | *LEU2 / leu2Δ HIS1 / his1Δ* | (Homann et al. 2009) |
| orf19.1253 | *pho4::LEU2 / pho4::HIS1* | (Homann et al. 2009) |
| orf19.4000 | *pho2::LEU2 / pho2::HIS1* | (Homann et al. 2009) |

#### C. glabrata strains

CG1 was generated from BG99 (Cormack and Falkow 1999) by inactivating the *URA3* gene with a random piece of DNA amplified from pUC19 plasmid and selecting on 5-FOA medium. CG1 was subsequently used to generate the rest of *C. glabrata* strains using homologous recombination with either antibiotic resistance or nutrient markers. The *NAT* gene conferring resistance to clonNat was amplified from the pFA6a-natNT2 plasmid in the PCR Toolbox collection (Janke et al. 2004). A standard S. *cerevisiae* transformation protocol was used with the following modification: instead of the short 40-60 bp flanking sequences, long flanking homologous sequences between 100-1000 bp were used on each end. The transformation constructs were generated using 2-step PCR with the split *URA3* marker (Reid, Lisby, and Rothstein 2002). ChIP-exo strains were made the same way as in S. *cerevisiae*, except using a C-terminal 3xFLAG tag (DYKDHDGDYKDHDIDYKDDDDK) for PHO4 with 6xG linker, and a N-terminal V5 tag (GKPIPNPLLGLDST) for PHO2 with 5xGS linker.

#### C. albicans strains

All *C. albicans* strains used in this study are from the transcriptional regulatory deletion library made by Homann et al (Homann et al. 2009) and ordered from the Fungal Genetics Stock Center (http://www.fgsc.net/)

### Media and Growth Conditions

Phosphate-free synthetic complete medium was prepared from Yeast Nitrogen Base with ammonium sulfate, without phosphates, without sodium chloride (MP Biomedicals, Santa Ana, California) and supplemented to a final concentration of 2% glucose, 1.5 mg/ml potassium chloride, 0.1 mg/ml sodium chloride and amino acids, as described previously (Lam, Steger, and O’Shea 2008). Monobasic potassium phosphate (1M solution, Sigma-Aldrich, St. Louis, MO) was added to phosphate-free medium to make high phosphate (Pi) medium containing a final concentration of 10 mM Pi. All media were adjusted to pH 4.0 with HCl. Yeast strains were grown at 30 °C with shaking and cell samples were collected at early/mid−logarithmic phase (OD_600_ 0.3 - 0.4). To induce the phosphate starvation response, yeast cells were first grown in 10 mM Pi medium to early/mid−logarithmic phase. Cells were then harvested by filtering and washed with 2-3 volumes of no Pi medium pre-warmed to 30 °C. Finally, cells were resuspended in pre-warmed no Pi medium and grown at 30 °C for 1 hour before being harvested for downstream analyses. For plate growth assays, yeast cells were grown in 10 mM Pi medium until mid−logarithmic phase, washed 2-3 times and re-suspended in sterile water. A 1:4 dilution series were made with sterile water in 96-well plates. A 48-pin tool was used to transfer ˜10 μL of resuspended cell culture onto appropriate solid agar plates. After 24-48 hours of growth at 30°C, pictures were taken using a standard gel box apparatus.

### Phosphatase assays

For the semi-quantitative assay, cells were grown overnight (preconditioning), diluted to OD_600_ ˜ 0.1 in the morning and grown to OD_600_ = 0.6-1. The cell culture was centrifuged, washed and resuspended in water. A four-fold serial dilution was prepared for each strain and spotted onto an agar plate using a 48-pin tool. Complete synthetic medium was used for *pho80Δ* strains and phosphate-free medium (see above) used for *PHO80* wild-type strains. After overnight growth, the agar plates were overlaid with Fast Blue Salt B stain (Sigma-Aldrich D9805), 1-naphthyl phosphate (1-NP, Sigma-Aldrich, D5602), and 1% agar in 0.1 m sodium acetate (pH 4.2) (Wykoff et al. 2007). Pictures were taken on a HP color scanner after 5 minutes.

For the quantitative phosphatase assay, cells were preconditioned and grown the same way as above. After collection by centrifugation, cells were washed and re-suspended with sterile water to OD_600_ ˜ 5. Then 30 μL of the re-suspended culture was transferred to a 96-well assay plate in triplicates. The cell re-suspension was incubated with 80 μL of 10 mM p-nitrophenyl phosphate (pNPP, Sigma-Aldrich P4744) dissolved in 0.1M sodium acetate (pH=4.2) for 15 min at 25°C. The reaction was quenched by adding 144 μL of saturated Na_2_CO_3_ (pH > 11) followed by 5 min of centrifugation at 3,000 g. Finally, 200 μL of the supernatant from each well was transferred to a new plate and OD_420_ was measured on a plate reader. Phosphatase activity was measured in units expressed as OD_420_/OD_600_ (Huang and O’Shea 2005).

### Transcriptional profiling by RNA-seq

All transcriptional profiling experiments were done in the *pho80Δ* background in both S. *cerevisiae* and *C. glabrata.* Two biological replicates (same genotype, but grown, collected and processed separately) were obtained for each sample. Briefly, yeast cells were collected using a cold methanol quenching method (Pieterse, Jellema, and van der Werf 2006; Zhou and O’Shea 2011). 20 mL of mid−log phase (OD_600_ = 0.2-0.5) cell cultures were added directly into 30 mL of pre-chilled methanol (˜ −50 °C), and incubated in an ethanol-dry ice bath at the same temperature for at least 20 minutes. Cells were collected by centrifugation and quickly washed with ice-cold water to remove methanol, and resuspended in RNAlater solution (Qiagen, Hilden, Germany). For each sample, 5*10^7^ cells were collected and mechanically lysed on a Minibeadbeater (BioSpec Products, Bartlesville, OK): Zirconia beads (0.5 mm, BioSpec Products #11079105z) were added to ˜600 μL of cell suspension per sample in a 2mL screw cap tube to the meniscus. Cells were lysed by four rounds of 1 min bead beating and 2 min of cooling in an ice-water bath. An RNasy Mini kit (Qiangen) was used to isolate total RNA from the lysed cell. RNA-seq libraries were prepared with the TruSeq RNA Library Preparation Kit v2 (Illumina, San Diego, CA) with the mRNA purification option, following the manufacturer’s protocol. The resulting libraries were sequenced on an Illumina HiSeq 2000, which produced on average 10 million 50 bp single end reads for each sample.

### Chromatin Immunoprecipitation

#### Biotin-tag Chromatin Immunoprecipitation (Bio-ChIP) in S. cerevisiae

Bio-ChIP was modified from techniques previously described (Zhou and O’Shea 2011; Lam, Steger, and O’Shea 2008; Kolodziej et al. 2009; van Werven and Timmers 2006), and performed for ScPho4 and CgPho4 in S. *cerevisiae* with the negative regulator ScPho80 deleted (*pho80*Δ). In addition, CgPho4 ChIP was also performed in S. *cerevisiae* with both ScPho80 and ScPho2 deleted (*pho80*Δ *pho2*Δ). One biological sample was analyzed for each strain. Two types of controls were included, i.e. an input sample, which is sonicated chromatin not subject to immune-precipitation (IP), and a mock sample, which uses a strain lacking the epitope tag recognized by the antibody, and is subject to the same IP as done for the biological sample. ˜100 OD_600_ units of cells were collected in phosphate-replete (10mM Pi) conditions. Cells were cross-linked with 1% formaldehyde (Fisher-Scientific #AC41073-1000, Hampton, NH) for 10 minutes and then quenched with 125 mM glycine for 10 minutes at room temperature. Cells were collected by centrifugation for 15 min at 6,000 g, immediately washed with cold PBS buffer (137 mM NaCl, 2.7 mM KCl, 10 mM Sodium phosphate dibasic, 2 mM potassium phosphate monobasic, pH 7.4), and mechanically lysed with 0.5 mm Zirconia beads in lysis buffer (50 mM HEPES, pH 7.5, 140 mM NaCl, 1 mM EDTA, 1% Triton X-100, 0.1% Na-Deoxycholate). Chromatin was fragmented by sonication using the Covaris E220 Adaptive Focus system (Covaris, Woburn, MA) using the 130 μL tube and the following setting: Duty Factor: 10%; Peak Incident Power: 175; Cycles/Burst: 200; Time:150s (Elfving et al. 2014). The cell lysate was then incubated with Dynabeads^®^ MyOne™ Streptavidin C1 (Invitrogen, Carlsbad, CA) overnight at 4°C. Afterwards, the dynabeads were washed with lysis buffer, high salt wash buffer (50 mM HEPES, pH 7.5, 500 mM NaCl, 1 mM EDTA, 1% Triton X-100, 0.1% Na-Deoxycholate), lithium wash buffer (10 mM Tris/HCl, pH 8.0, 500 mM LiCl, 1 mM EDTA, 1% NP-40, 0.1% Na-Deoxycholate), and SDS wash buffer (10 mM Tris/HCl, pH 8.0, 1 mM EDTA, 3% SDS) for 2 × 2 minutes at room temperature, and 1 × 2 minutes with TE buffer (10 mM Tris/HCl, pH 8.0, 1 mM EDTA). Crosslinking was reversed by incubation of samples at 65 °C for at least 6 hours in 10 mM Tris-HCl, pH 8.0, 1 mM EDTA, 0.8% SDS. RNA and proteins in the samples were digested with 20 μg/ml RNase A (Thermo Fisher Scientific, Waltham, MA) at 37 °C for 2 hours and 0.2 mg/ml proteinase K (Roche, Basel, Switzerland) for 2 hours at 55 °C. DNA was then purified with minElute column (Qiagen). 5% of the volume of cell lysate was removed after sonication and used to prepare the input DNA for each ChIP experiment. ChIP libraries were prepared using the NEBNext^®^ Ultra™ II DNA Library Prep Kit (New England Laboratory, Ipswich, MA) and sequenced on an Illumina HiSeq2000 instrument to produce ˜10 million 50 bp single end reads per sample.

#### Chromatin Immunoprécipitation exonuclease (ChIP-exo) in C. glabrata

ChIP-exo was performed in *C. glabrata* for C-3xFLAG tagged CgPho4 either with or without CgPho2, and N-V5 tagged CgPho2 following published methods (Rhee and Pugh 2012; Serandour et al. 2013). All strains have wild-type CgPho80. Only mock samples, but not input samples, were included because the latter didn’t apply to the ChIP-exo procedure (Rhee and Pugh 2012). One biological sample was analyzed for each strain. Briefly, cells were fixed and processed in the same way as in Bio-ChIP until the sonication step. Afterwards, either the AntiFLAG^®^ M2 Magnetic Beads (Sigma-Aldrich #M8823) or the Anti-V5 antibody (Abcam #15828, Cambridge, UK) was incubated with the cell lysates for 2 hours up to overnight. For Anti-V5 antibody, Dynabeads™ Protein G (Thermo Fisher Scientific #10003D) was added to the antibody-lysate mix and incubated for an additional 2 hours at 4°C. The following steps are modified from the Active Motif ChIP-exo vA4 manual (http://www.activemotif.com/documents/1938.pdf): with the chromatin still bound to the beads, the DNA was end-polished and P7-exo adapters are ligated onto the blunt ends. The nicked DNA was repaired and then digested by lambda and RecJf exonulceases to excise DNA in a 5’ to 3’ direction, trimming up to the site of the cross-linking and selectively eliminating the P7 adapter at the 5 end. Following cross-link reversal and elution from the beads, the DNA was made double-stranded by P7 primer extension and a P5-exo adapter was added to the exonuclease-treated ends. The DNA library was PCR amplified and size selected before it was subjected to high-throughput sequencing.

### Functional genomics data analysis

#### RNA-seq: differential expression and mutant cycle analysis

##### Reads mapping and counting

The raw reads were mapped to the corresponding species reference genomes (S. *cerevisiae* genome version S288C_R64-1-1; *C. glabrata* genome version s02-m02-r09) using Bowtie (v1.1.1) with the option ‘-m 1 --best-strata’ (Langmead et al. 2009). The resulting SAM files were sorted using Samtools (v1.2) (Li et al. 2009) and the number of reads per transcript was counted using Bedtools2 (Quinlan and Hall 2010), with the option ‘bedtools coverage -a BAM_file -b genome_annotation.bed -S -s sorted -g Chrom.length’. Gene features for S. *cerevisiae* were downloaded from the Saccharomyces Genome Database (SGD), version R64- 2-1; gene features for *C. glabrata* were downloaded from the Candida Genome Database (CGD), version s02-m07-r04. The count matrix was imported into R for downstream analyses.

##### Differential gene expression analysis in S. *cerevisiae*

First, 676 genes having <1 count per million reads in at least 3 samples were removed, leaving 5707 genes in the dataset. Next, we used the trimmed mean of M-values ("TMM") method in the EdgeR package to calculate the normalization factors for scaling the raw library sizes (Robinson and Oshlack 2010; Robinson, McCarthy, and Smyth 2010), and applied voom transformation (Law et al. 2014) to remove the mean-variance relationship on the log_2_ transformed count data.

To identify genes induced by each Pho4 ortholog in the S. *cerevisiae* background, we compared the transcriptional profiles between a strain with the Pho4 ortholog and a strain without Pho4, either with or without *ScPHO2*, e.g. *CgPHO4 ScPHO2* vs *CgPHO4 pho2*Δ. Two biological replicates were analyzed for each strain. With eight Pho4 orthologs, there are a total of 16 pairwise comparisons for each gene in the dataset. In practice, we analyzed all 16 comparisons with a single linear model framework using the LIMMA package in R (Ritchie et al. 2015). The normalized log2 read count for gene *i*, strain *j* and replicate *k* can be expressed as ^*£i,j,k*^, where *j* = {*pho4*Δ.*pho2*Δ, *pho4*Δ.*ScPHO2, PHO4*.pho2*Δ, *PHO4*.ScPHO2*}, with *PHO4\** replaced by one of the eight Pho4 orthologs, and *k* = 1, 2. The fold induction for gene *i* by ScPho4 in the presence of ScPho2 can then be estimated as *β*_*i,ScPHO*4.*ScPHO*2_ – *β*_*i,pho4*Δ.*ScPH02*_. To test for significant differences in the expression of gene *i* between two genotypes, LIMMA performs a moderated t-test, which differs from a standard t-test in that it uses an Empirical Bayes method to moderate the standard error term for each gene so as to make the estimates more robust (Smyth, Michaud, and Scott 2005). After obtaining the 16 fold induction estimates for each gene (8 Pho4 orthologs with or without ScPho2), we tested for significant induction against a null hypothesis of *fold change <= 2*, with the function treat(…, lfc=1). The raw p-values were then *pooled* across all genes and all pairwise comparisons, and a Benjamini-Hochberg procedure was used to control the false discovery rate at 0.05, with the function decideTests(…, p.value = 0.05, method = “global-) in LIMMA. The advantage of this approach is that the raw p-value cutoff is consistent across all 16 comparisons.

The estimated fold changes were used to produce Figure 3A and 3B, for the 247 genes that were significantly induced more than two-fold by at least one of the Pho4 orthologs. For Figure 3C, we calculated the ratios between the fold induction without vs with ScPho2 for each Pho4 ortholog, for the 16 genes significantly induced by all eight Pho4 orthologs. For Figure 4C, the number of genes significantly induced by each Pho4 ortholog with ScPho2 was determined by the method described above.

##### Mutant cycle analysis in *C. glabrata*

In order to explicitly evaluate the contribution from CgPho4, CgPho2 and their collaboration to gene induction in *C. glabrata*, we applied the mutant cycle analysis method (Capaldi et al. 2008; Zhou and O’Shea 2011), which decomposes the induction fold change of each gene into Pho4 acting alone (Pho4), Pho2 acting alone (Pho2) and Pho4/Pho2 collaborative effect (CO). To estimate the three components, we used the same linear model approach as in S. *cerevisiae*, where the fold induction for any gene *i* in strain *j* and replicate *k* is expressed as *y_i,j,k_* = *β_i,j_* + *£_i,j,k_*, where *j* = {*CgPHO4.CgPHO2, CgPHO4.pho2*Δ, *pho4*Δ.*CgPHO2, pho4*Δ.*pho2*Δ}, k = 1,2 and *£_i,j,k_* represents the noise. The three expression components can then be estimated as

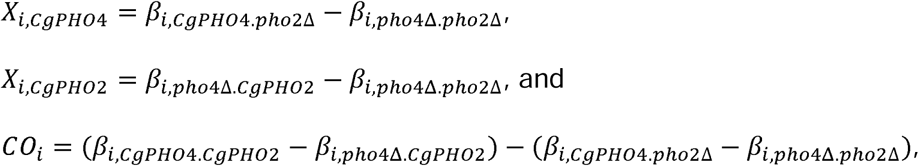

We performed this analysis for the 79 genes that are directly bound and induced by CgPho4 in *C. glabrata* (Figure 5- source data 2). The genes were then hierarchically clustered using Ward’s method (Ward 1963; Murtagh and Legendre 2014) based on the estimates of the three expression components.

##### ChIP-seq analysis

We combined automatic peak calling with manual curation in order to identify binding sites for ScPho4 and CgPho4 in S. *cerevisiae*, and CgPho4 and CgPho2 binding sites in *C. glabrata.* First, we mapped the Bio-ChIP and ChIP-exo sequencing data to the respective reference genomes, as we did for RNA-seq data. Second, we used the program GEM (v2.6) (Guo, Mahony, and Gifford 2012) to automatically call peaks for ScPho4 and CgPho4 ("--smooth 30 -fold 1.5") with mock sample as control. GEM iteratively performs peak calling and *de novo* motif discovery to improve both sensitivity and specificity (Guo, Mahony, and Gifford 2012) – for CgPho4, GEM identified the same E-box motif “CACGTG- as recognized by ScPho4 (Zhou and O’Shea 2011) in both S. *cerevisiae* and *C. glabrata.* For CgPho2, however, GEM performed poorly, identifying <10 significant peaks and no significantly enriched motifs. Instead, we used MACS2 (Zhang et al. 2008) to call peaks for CgPho2 in *C. glabrata* with “-s 50 --bw 200 -q 0.01 --keep-dup auto --slocal 1000 -B --verbose 4 -m 2 100 --call-summits-, and identified 640 binding peaks with a fold enrichment (FE) greater than 3 over the mock sample (380 peaks with FE > 4 and 1565 peaks with FE > 2). *De novo* motif discovery using RSAT Peak-motifs algorithm (Thomas-Chollier et al. 2011) identified “CACAGA- as the top motif, which appears in 126 of the 380 (33.16%) peaks that have FE > 4. It bears no similarity with the *in vitro* identified ScPho2 motif ("ATTA") (Zhu et al. 2009; Zhao, Granas, and Stormo 2009). Nor did it match any known motifs of yeast transcription factors (de Boer and Hughes 2012). In subsequent analyses, we used CgPho2 ChIP peaks for categorizing CgPho4 binding sites into ones cobound by CgPho2 or those bound by CgPho4 only.

After the automatic peak calling, we manually curated the peaks with visual help from a genome browser tool, MochiView (Homann and Johnson 2010). We calculated a fold enrichment trace for each ChIP sample by comparing it to its matching input (or mock, in case of ChIP-exo), using MACS2 following http://github.com/taoliu/MACS/wiki/Build-Signal-Track. This trace file is converted to WIG format (http://genome.ucsc.edu/goldenPath/help/wiggle.html) and loaded into MochiView, along with the coordinates of the GEM identified ChIP peaks. We also imported previously published datasets for ScPho4 occupancy in S. *cerevisiae* with or without ScPho2 in no Pi conditions (Zhou and O’Shea 2011). We then examined each GEM identified ChIP peak, using the height and peak shape to identify spurious binding peaks and false negative ones. The curated datasets are presented as source data accompanying Figure 4 and Figure 5.

To estimate the ratio of ScPho4 and CgPho4 occupancy without vs with Pho2 in S. *cerevisiae* (Figure 4B), we combined the published ScPho4 binding occupancy in no phosphate conditions (Zhou and O’Shea 2011) with our data for CgPho4 in the *pho80Δ* background. For the former, we downloaded the WIG files from the NCBI GEO database for series GSE29506, sample GSM730517 (Pho4_ChIP_NoPi) and GSM730527 (Pho4_ChIP_dPho2_NoPi), and scaled the raw library sizes so that the average coverage per base pair is 1. Then these were analyzed along with our CgPho4 ChIP enrichment in *pho80*Δ. For each of the ScPho4 (74) and CgPho4 (115) ChIP peaks, we recorded the maximum value of the normalized read counts for ScPho4 and CgPho4, with or without Pho2, and calculated the ratio between the two. Then the −log_2_ transformed values were plotted (Figure 4B). A two-sided Student’s t-test was performed to assess the significance of the difference in the group means.

To investigate the nucleosome and Cbf1 occupancy in S. *cerevisiae* at sites either bound by CgPho4 only or by both ScPho4 and CgPho4, we used the published data from Table S2 of (Zhou and O’Shea 2011). We first translated the coordinates in the table from sacCer1 into sacCer3 using the Galaxy tool (Afgan et al. 2016). We then identified all “CACGTG- motifs overlapping with the ChIP peaks for ScPho4 or CgPho4 identified in this study using GenomicsRanges package in R (Lawrence et al. 2013), and extracted the corresponding “Nucleosomes.High.Pi- and “Cbf1.Enrichment.High.Pi- values from the table. Finally, the occupancy or enrichment are plotted separately for sites bound by both ScPho4 and CgPho4, and sites bound by only CgPho4. A two-sided Student’s t-test was performed to assess the significance of the difference in the group means.

##### Identification of CgPho4 targets in S. cerevisiae and C. glabrata

To identify ScPho4 and CgPho4 induced genes in S. *cerevisiae* and CgPho4 induced genes in *C. glabrata*, we assigned each ChIP peak to the nearest gene(s) on both sides, provided that the ChIP peak is upstream (in the 5’ UTR) of the gene, and queried the gene induction results from the RNA-seq experiments. ChIP binding peaks associated with Pho4-dependent gene induction are identified as the direct targets of ScPho4 or CgPho4. The identified target genes are presented in the source data accompanying Figure 4 and Figure 5.

#### Annotating Pho4 targets in *C. glabrata*

Annotation of putative Pho4 targets in *C. glabrata* was based largely on information of the orthologs in the well-annotated S. *cerevisiae* and *C. albicans* genomes. Gene Ontology enrichment analysis was performed using the “GO term finder- tool, and mapping genes to GO Slim terms performed using the “GO Slim mapper-, both available on the CGD website (http://www.candidagenome.org/). All genomic features were used as the background set. For the enrichment analysis, a threshold of *P* < 0.1 was used for selecting significant GO terms. We focused on the “Biological Processes- to identify the functional groups as reported in the results.

## Acknowledgements

We thank Cecile Fairhead, Rodney Rothstein, Aviv Regev and Suzanne Noble for sharing yeast and bacteria strains. Cecile Fairhead shared Pho4 and Pho2 ortholog sequences in the newly sequenced *glabrata* clade species. We thank Elmar Czeko for help with ChIP-exo. We thank Marty Kreitman, Christopher Chidley, Joseph Piechura, Xiaoyu Zheng, Alex Nyugen-Ba, Dennis Wykoff, Andrew Murray, Julia Koehler and members of the O’Shea lab for discussion and critical reading of the manuscript. Illumina sequencing for RNA-seq and ChIP-seq was performed at the Bauer Core Facility, and the computational analysis done on the Odyssey cluster, both supported by the FAS Division of Science at Harvard University. The Howard Hughes Medical Institute supported this work.

---

**Figure 3- figure supplement 1.**
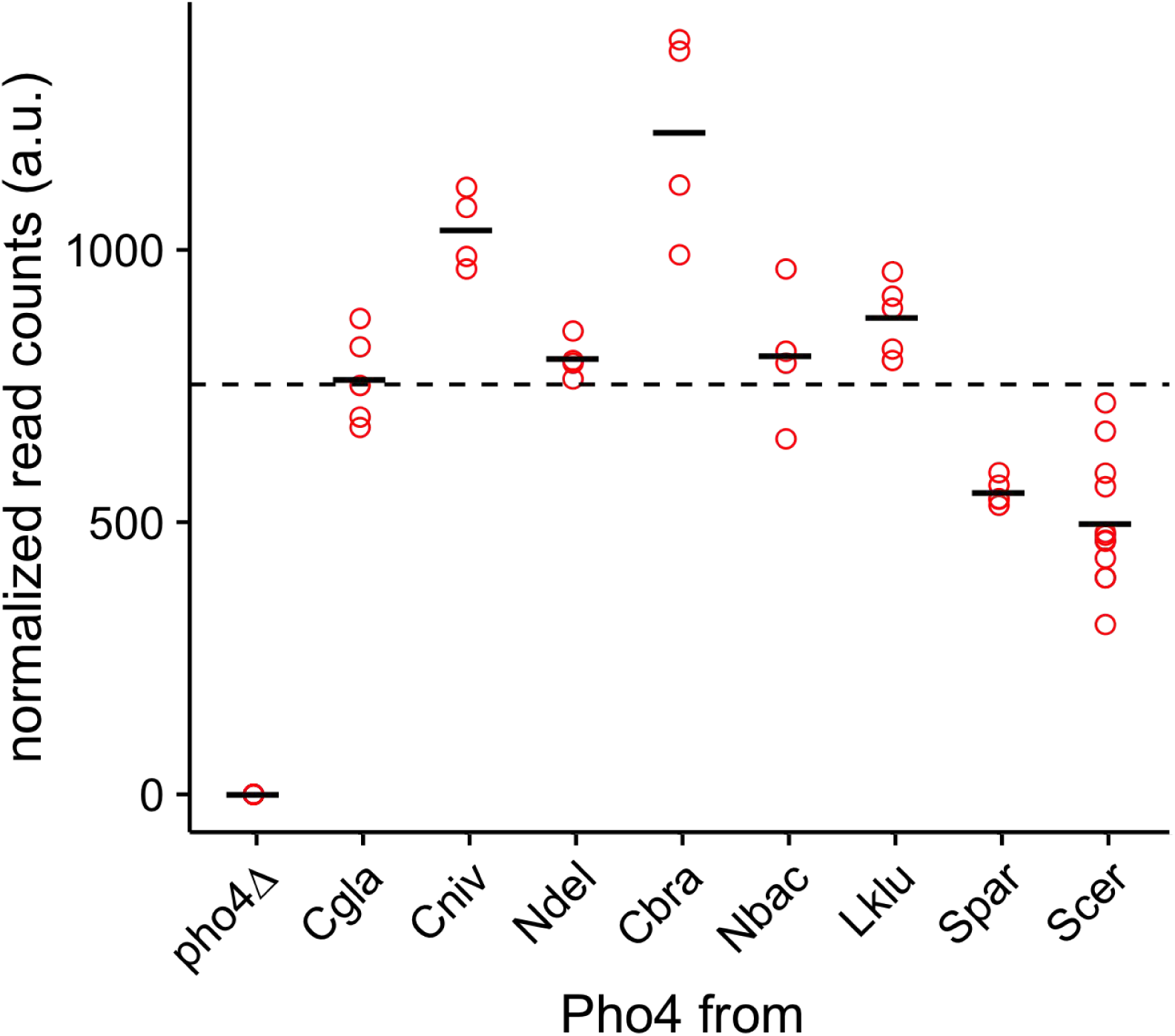
Expression levels of the Pho4 orthologs by RNA-seq. The scatter plot shows normalized transcript abundance for the *PHO4* orthologs. The black bar indicates the mean for each *PHO4* ortholog. The dotted line shows the average expression level over all Pho4 orthologs (except *pho4Δ*).

**Figure 3- figure supplement 2.**
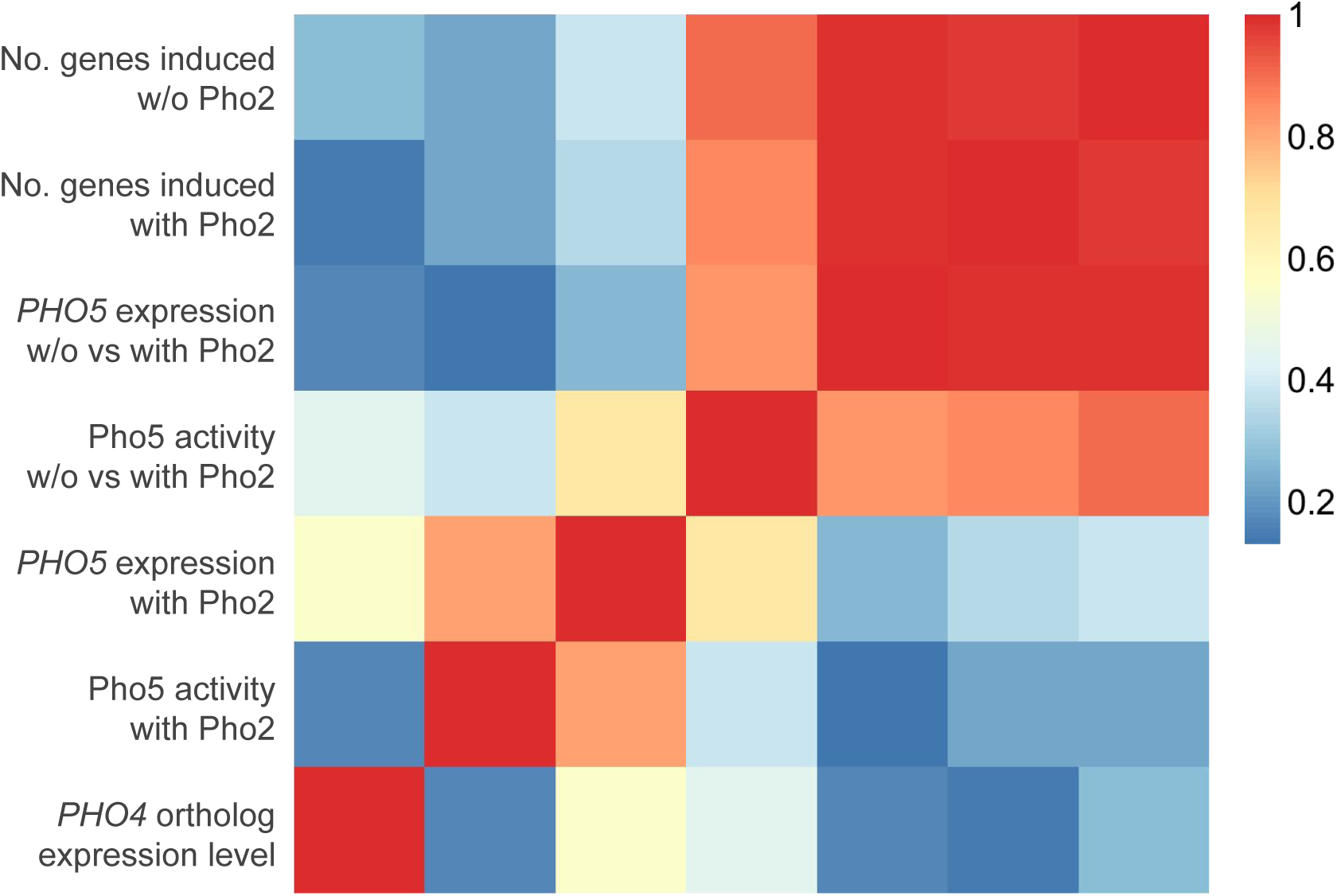
Expression level of a Pho4 ortholog does not correlate with its level of Pho2-dependence or the number of genes it induces. The heatmap shows the Pearson correlation between a Pho4 ortholog’s expression level and the corresponding strain’s gene induction statistics, including a) phosphatase assay activity; b) *PHO5* expression level with Pho2; c) ratio of phosphatase assay activity without vs with Pho2; d) ratio of *PHO5* expression level without vs with Pho2; e) number of genes induced by each Pho4 ortholog with Pho2 and f) number of genes induced by each Pho4 ortholog without Pho2.

**Figure 3- source data 1.** This zip file contains four tab-delimited csv files. Two of them (fold_with_Pho2.csv and fold_no_pho2.csv) record the gene fold change estimates for each Pho4 ortholog with or without ScPho2. The other two (sig_with_Pho2.csv and sig_no_pho2.csv) record for each Pho4 ortholog whether a gene is deemed significantly induced with or without Pho2.

**Figure 4- figure supplement 1.**
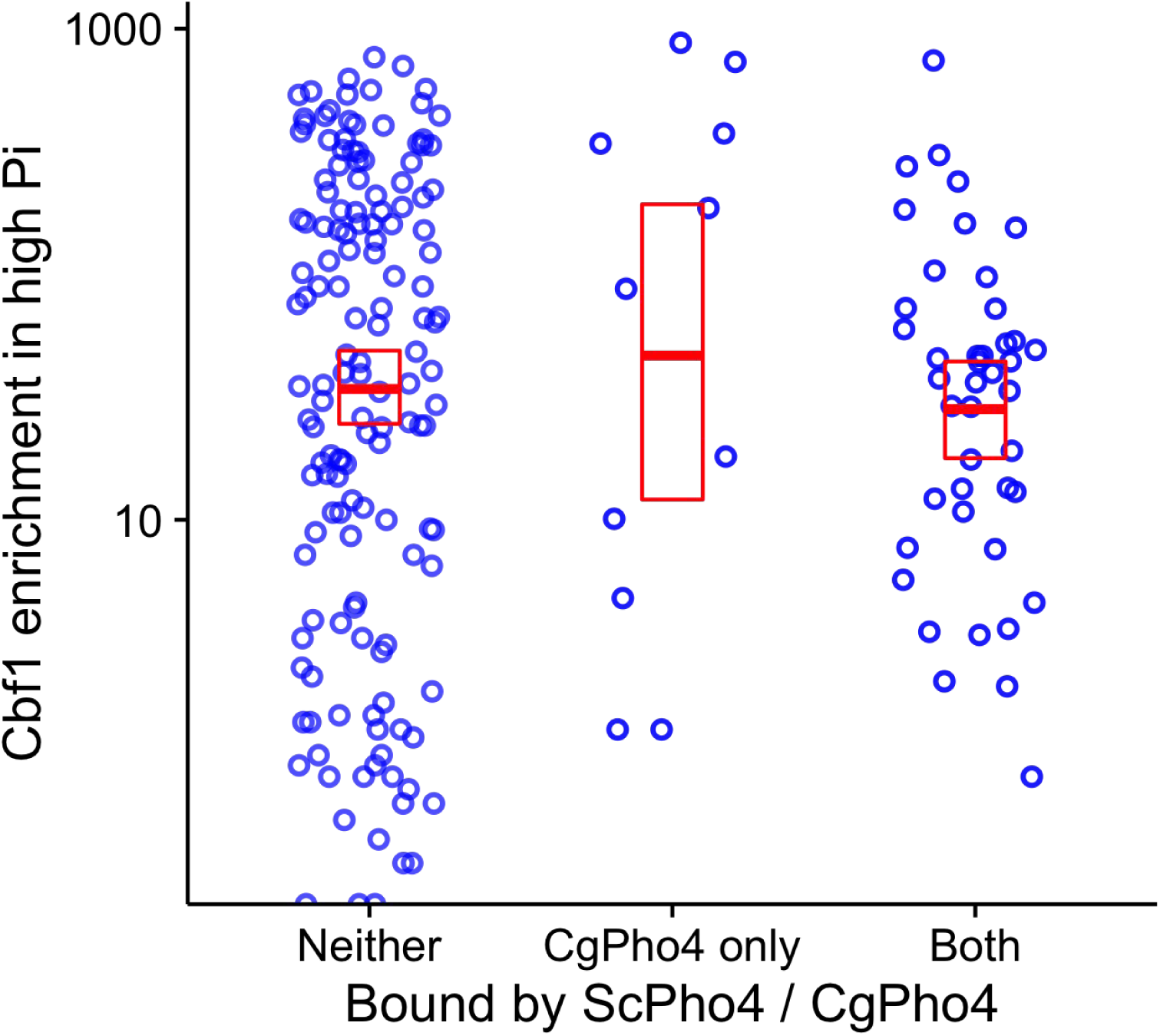
Cbf1 enrichment in high Pi conditions in *S. cerevisiae* is not significantly different between sites bound by both CgPho4 and ScPho4 and sites bound by CgPho4 only. Scatter plot shows the log_10_ transformed Cbf1 enrichment in high Pi conditions (Zhou and O’Shea 2011) at a subset of the E-box motifs*, grouped by whether the motif is bound by neither ScPho4 and CgPho4, only CgPho4 or both. A two-sided t-test comparing the means of the latter two groups yielded a *p*-value of 0.53. * To avoid the confounding nucleosome exclusion, the E-box motifs in the S. *cerevisiae* genome were ordered by their nucleosome occupancy in high Pi conditions (Zhou and O’Shea 2011) from low to high, and the top 25% most accessible ones were used for this plot.

**Figure 4- source data 1. List of ChIP-identified binding sites of ScPho4 and CgPho4 in** *S. cerevisiae*, **and associated gene information.** Yellow highlight of chromosome names and coordinates mark the same ChIP peak associated with two downstream genes. Two genes (YOR183W and YNL042W-B) do not have induction fold change estimates because of low or no coverage in the RNA-seq data, and marked “NA-. YDL106C *(PHO2)* is deleted in a subset of the strains and therefore excluded from the differential gene expression analysis. Five genes (in red) are exceptions in that the associated CgPho4 ChIP peaks (none of them are bound by ScPho4) were in their coding sequences. For two of the five genes, i.e. *RIM15* and YDR089W, CgPho4 binding is associated with CgPho4-dependent gene induction.

**Figure 5- figure supplement 1.**
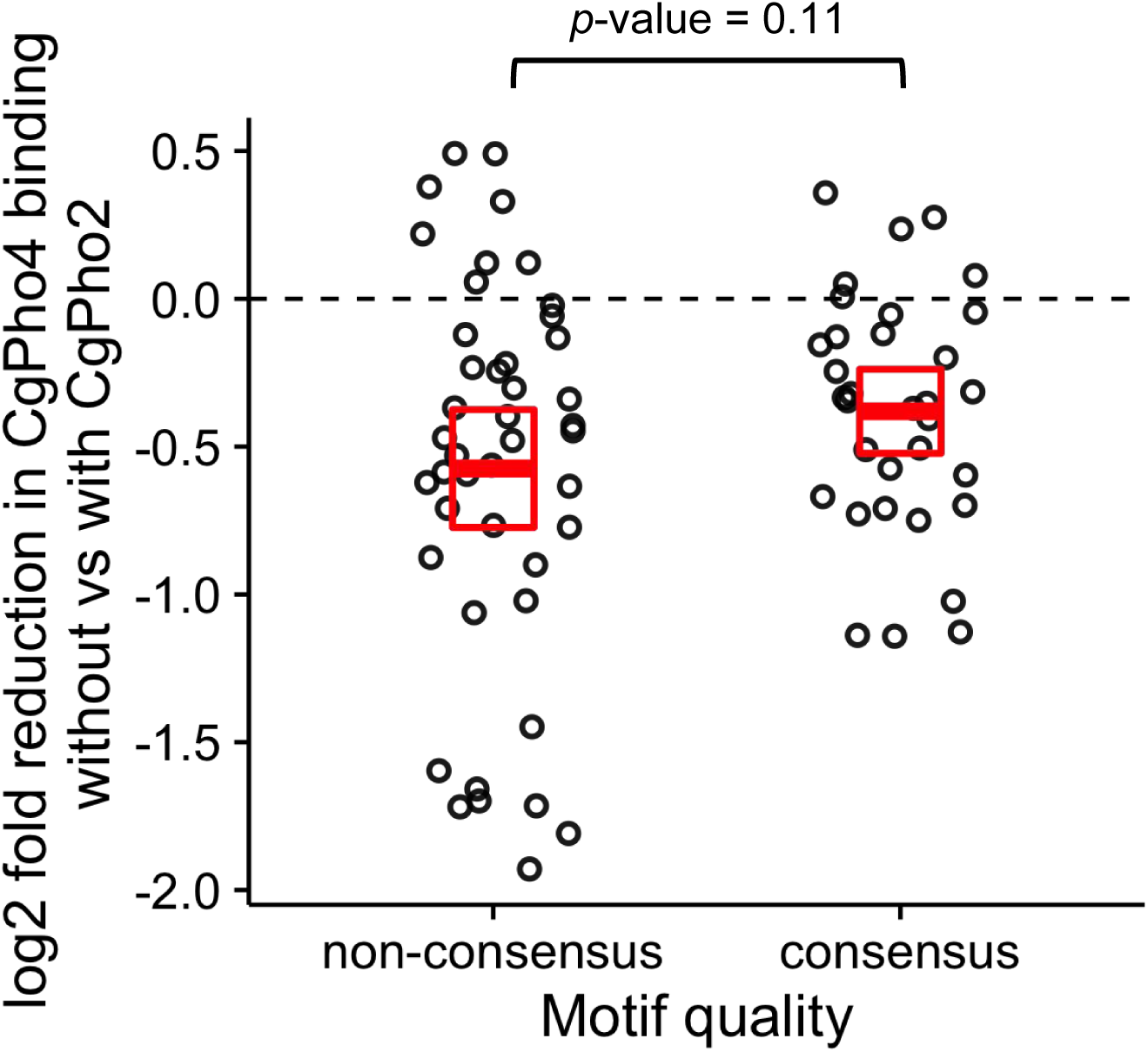
Comparison of CgPho2 influence on CgPho4 binding at sites with and without the consensus motif. Scatter plot showing log_2_ ratio of CgPho4 ChIP enrichment without vs with CgPho2. The thick red bar depicts the mean and the box the 95% confidence limits computed by non-parametric bootstrapping. The difference in the means of the two groups is not significant at a 0.05 level by a Student’s t test, with a *p*-value = 0.11.

**Figure 5- source data 1. List of ChIP-identified binding sites of CgPho4 in** *C.glabrata.*

**Figure 5- source data 2. List of CgPho4 directly bound and induced genes and the associated expression components from the mutant cycle analysis.**

**Figure 6- source data 1. Gene Ontology terms enrichment analysis results**

**Figure 6- source data 2. Table comparing ScPho4 and CgPho4 targets with phosphate homeostasis related functions**

**Figure 6- source data 3. All 79 CgPho4 targets in** *C. glabrata* **mapped to Gene Ontology Slim terms**

**Figure 6- source data 4. Contains five tables listing CgPho4 target genes annotation grouped by functional categories: Table S1- Non-phosphate related stress and starvation response; Table S2 - Response to chemicals; Table S3 - Cell wall and cell adhesion; Table S4 - Carbohydrate metabolism; Table S5 - all other functional groups.**

## References

Afgan, Enis, Dannon Baker, Marius van den Beek, Daniel Blankenberg, Dave Bouvier, Martin Čech, John Chilton, et al. 2016. “The Galaxy Platform for Accessible, Reproducible and Collaborative Biomedical Analyses: 2016 Update.- Nucleic Acids Research 44 (W1): W3–10. doi:10.1093/nar/gkw343.

Ang, Yen-Sin, Renee N. Rivas, Alexandre J. S. Ribeiro, Rohith Srivas, Janell Rivera, Nicole R. Stone, Karishma Pratt, et al. 2016. “Disease Model of GATA4 Mutation Reveals Transcription Factor Cooperativity in Human Cardiogenesis.- Cell 167 (7): 1734–1749. e22. doi:10.1016/j.cell.2016.11.033.

Atanasova, Ralitsa, Adela Angoulvant, Maurel Tefit, Frédérick Gay, Juliette Guitard, Dominique Mazier, Cécile Fairhead, and Christophe Hennequin. 2013. “A Mouse Model for Candida Glabrata Hematogenous Disseminated Infection Starting from the Gut: Evaluation of Strains with Different Adhesion Properties.- PloS One 8 (7): e69664. doi:10.1371/journal.pone.0069664.

Barbarić, S., M. Münsterkötter, J. Svaren, and W. Hörz. 1996. “The Homeodomain Protein Pho2 and the Basic-Helix-Loop-Helix Protein Pho4 Bind DNA Cooperatively at the Yeast PHO5 Promoter.- Nucleic Acids Research 24 (22): 4479–86.

Barbaric, Slobodan, Martin Münsterkötter, Colin Goding, and Wolfram Hörz. 1998. “Cooperative Pho2-Pho4 Interactions at thePHO5 Promoter Are Critical for Binding of Pho4 to UASp1 and for Efficient Transactivation by Pho4 at UASp2.- Molecular and Cellular Biology 18 (5): 2629–39.

Boer, Carl G. de, and Timothy R. Hughes. 2012. “YeTFaSCo: A Database of Evaluated Yeast Transcription Factor Sequence Specificities.- Nucleic Acids Research 40 (Database issue): D169–179. doi:10.1093/nar/gkr993.

Byrne, Kevin P., and Kenneth H. Wolfe. 2005. “The Yeast Gene Order Browser: Combining Curated Homology and Syntenic Context Reveals Gene Fate in Polyploid Species.- Genome Research 15 (10): 1456–61. doi:10.1101/gr.3672305.

Capaldi, Andrew P., Tommy Kaplan, Ying Liu, Naomi Habib, Aviv Regev, Nir Friedman, and Erin K. O’Shea. 2008. “Structure and Function of a Transcriptional Network Activated by the MAPK Hog1.- Nature Genetics 40 (11): 1300–1306. doi:10.1038/ng.235.

Cheatle Jarvela, Alys M., Lisa Brubaker, Anastasia Vedenko, Anisha Gupta, Bruce A. Armitage, Martha L. Bulyk, and Veronica F. Hinman. 2014. “Modular Evolution of DNA-Binding Preference of a Tbrain Transcription Factor Provides a Mechanism for Modifying Gene Regulatory Networks.- Molecular Biology and Evolution 31 (10): 2672–88. doi:10.1093/molbev/msu213.

Cormack, B. P., and S. Falkow. 1999. “Efficient Homologous and Illegitimate Recombination in the Opportunistic Yeast Pathogen Candida Glabrata.- Genetics 151 (3): 979–87.

De Las Penas, Alejandro, Jacqueline Juárez-Cepeda, Eunice López-Fuentes, Marcela Briones-Martín-Del-Campo, Guadalupe Gutiérrez-Escobedo, and Irene Castaño. 2015. “Local and Regional Chromatin Silencing in Candida Glabrata: Consequences for Adhesion and the Response to Stress.- FEMS Yeast Research 15 (6). doi:10.1093/femsyr/fov056.

Diezmann, Stephanie, Cymon J. Cox, Gabriele Schönian, Rytas J. Vilgalys, and Thomas G. Mitchell. 2004. “Phylogeny and Evolution of Medical Species of Candida and Related Taxa: A Multigenic Analysis.- Journal of Clinical Microbiology 42 (12): 5624–35. doi:10.1128/JCM.42.12.5624-5635.2004.

Domergue, Renee, Irene Castaño, Alejandro De Las Peñas, Margaret Zupancic, Virginia Lockatell, J. Richard Hebel, David Johnson, and Brendan P. Cormack. 2005. “Nicotinic Acid Limitation Regulates Silencing of Candida Adhesins During UTI.- Science 308 (5723): 866–70. doi:10.1126/science.1108640.

Elfving, Nils, Rãzvan V. Chereji, Vasudha Bharatula, Stefan Björklund, Alexandre V. Morozov, and James R. Broach. 2014. “A Dynamic Interplay of Nucleosome and Msn2 Binding Regulates Kinetics of Gene Activation and Repression Following Stress.- Nucleic Acids Research 42 (9): 5468–82. doi:10.1093/nar/gku176.

Fabre, Emeline, Ghenima Sfihi-Loualia, Marilyne Pourcelot, Bernadette Coddeville, Frédéric Krzewinski, Julie Bouckaert, Emmanuel Maes, et al. 2014. “Characterization of the Recombinant Candida Albicans β-1,2-Mannosyltransferase That Initiates the ß-Mannosylation of Cell Wall Phosphopeptidomannan.- Biochemical Journal 457 (2): 347–60. doi:10.1042/BJ20131012.

Gabaldón, Toni, Tiphaine Martin, Marina Marcet-Houben, Pascal Durrens, Monique Bolotin-Fukuhara, Olivier Lespinet, Sylvie Arnaise, et al. 2013. “Comparative Genomics of Emerging Pathogens in the Candida Glabrata Clade.- BMC Genomics 14 (1): 623. doi:10.1186/1471-2164-14-623.

Gietz R. Daniel, and Robert H. Schiestl. 2007. “High-Efficiency Yeast Transformation Using the LiAc/SS Carrier DNA/PEG Method.- Nature Protocols 2 (1): 31–34. doi:10.1038/nprot.2007.13.

Gompel, Nicolas, Benjamin Prud’homme, Patricia J. Wittkopp, Victoria A. Kassner, and Sean B. Carroll. 2005. “Chance Caught on the Wing: Cis-Regulatory Evolution and the Origin of Pigment Patterns in Drosophila.- Nature 433 (7025): 481–87. doi:10.1038/nature03235.

Guo, Yuchun, Shaun Mahony, and David K. Gifford. 2012. “High Resolution Genome Wide Binding Event Finding and Motif Discovery Reveals Transcription Factor Spatial Binding Constraints.- PLOS Comput Biol 8 (8): e1002638. doi:10.1371/journal.pcbi.1002638.

Harrell, Frank E Jr. 2016. Hmisc: Harrell Miscellaneous (version 3.17-2). R. http://CRAN.R-project.org/package=Hmisc.

Homann, Oliver R., Jeanselle Dea, Suzanne M. Noble, and Alexander D. Johnson. 2009. “A Phenotypic Profile of the Candida Albicans Regulatory Network.- PLoS Genet 5 (12): e1000783. doi:10.1371/journal.pgen.1000783.

Homann, Oliver R., and Alexander D. Johnson. 2010. “MochiView: Versatile Software for Genome Browsing and DNA Motif Analysis.- BMC Biology 8: 49. doi:10.1186/1741-7007-8-49.

Huang, Sidong, and Erin K. O’ Shea. 2005. “A Systematic High-Throughput Screen of a Yeast Deletion Collection for Mutants Defective in PHO5 Regulation.- Genetics 169 (4): 1859–71. doi:10.1534/genetics.104.038695.

Ihmels, Jan, Sven Bergmann, Maryam Gerami-Nejad, Itai Yanai, Mark McClellan, Judith Berman, and Naama Barkai. 2005. “Rewiring of the Yeast Transcriptional Network through the Evolution of Motif Usage.- Science (New York, N.Y.) 309 (5736): 938–40. doi:10.1126/science.1113833.

Ikeh, Mélanie A. C., Stavroula L. Kastora, Alison M. Day, Carmen M. Herrero-de-Dios, Emma Tarrant, Kevin J. Waldron, A. Peter Banks, et al. 2016. “Pho4 Mediates Phosphate Acquisition in Candida Albicans and Is Vital for Stress Resistance and Metal Homeostasis.- Molecular Biology of the Cell, July, mbc.E16-05-0266. doi:10.1091/mbc.E16-05-0266.

Janke, Carsten, Maria M. Magiera, Nicole Rathfelder, Christof Taxis, Simone Reber, Hiromi Maekawa, Alexandra Moreno-Borchart, et al. 2004. “A Versatile Toolbox for PCR-Based Tagging of Yeast Genes: New Fluorescent Proteins, More Markers and Promoter Substitution Cassettes.- Yeast 21 (11): 947–62. doi:10.1002/yea.1142.

Jawhara, Samir, Estelle Mogensen, François Maggiotto, Chantal Fradin, Aurore Sarazin, Laurent Dubuquoy, Emmanuel Maes, Yann Guérardel, Guilhem Janbon, and Daniel Poulain. 2012. “Murine Model of Dextran Sulfate Sodium-Induced Colitis Reveals Candida Glabrata Virulence and Contribution of ß-Mannosyltransferases.- The Journal of Biological Chemistry 287 (14): 11313–24. doi:10.1074/jbc.M111.329300.

Kaffman, A., I. Herskowitz, R. Tjian, and E. K. O’Shea. 1994. “Phosphorylation of the Transcription Factor PHO4 by a Cyclin-CDK Complex, PHO80-PHO85.- Science (New York, N.Y.) 263 (5150): 1153–56.

Kasper, Lydia, Katja Seider, and Bernhard Hube. 2015. “Intracellular Survival of Candida Glabrata in Macrophages: Immune Evasion and Persistence.- FEMS Yeast Research, June. doi:10.1093/femsyr/fov042.

Kerwin, Christine L., and Dennis D. Wykoff. 2009. “Candida Glabrata PHO4 Is Necessary and Sufficient for Pho2-Independent Transcription of Phosphate Starvation Genes.- Genetics 182 (2): 471–79. doi:10.1534/genetics.109.101063.

Kerwin, Christine L., and Dennis D. Wykoff. 2012. “De Novo Generation of a Phosphate Starvation-Regulated Promoter in Candida Glabrata.- FEMS Yeast Research 12 (8): 980–989. doi:10.1111/1567-1364.12000.

Knop, Michael. 2006. “Evolution of the Hemiascomycete Yeasts: On Life Styles and the Importance of Inbreeding.- BioEssays 28 (7): 696–708. doi:10.1002/bies.20435.

Kolodziej, Katarzyna E., Farzin Pourfarzad, Ernie de Boer, Sanja Krpic, Frank Grosveld, and John Strouboulis. 2009. “Optimal Use of Tandem Biotin and V5 Tags in ChIP Assays.- BMC Molecular Biology 10 (1): 6. doi:10.1186/1471-2199-10-6.

Komeili, Arash, and Erin K. O’Shea. 1999. “Roles of Phosphorylation Sites in Regulating Activity of the Transcription Factor Pho4.- Science 284 (5416): 977–80. doi:10.1126/science.284.5416.977.

Lam, Felix H., David J. Steger, and Erin K. O’Shea. 2008. “Chromatin Decouples Promoter Threshold from Dynamic Range.- Nature 453 (7192): 246–50. doi:10.1038/nature06867.

Langmead, Ben, Cole Trapnell, Mihai Pop, and Steven L. Salzberg. 2009. “Ultrafast and Memory-Efficient Alignment of Short DNA Sequences to the Human Genome.- Genome Biology 10: R25. doi:10.1186/gb-2009-10-3-r25.

Law, Charity W., Yunshun Chen, Wei Shi, and Gordon K. Smyth. 2014. “Voom: Precision Weights Unlock Linear Model Analysis Tools for RNA-Seq Read Counts.- Genome Biology 15: R29. doi:10.1186/gb-2014-15-2-r29.

Lawrence, Michael, Wolfgang Huber, Hervé Pagès, Patrick Aboyoun, Marc Carlson, Robert Gentleman, Martin T. Morgan, and Vincent J. Carey. 2013. “Software for Computing and Annotating Genomic Ranges.- PLOS Computational Biology 9 (8): e1003118. doi:10.1371/journal.pcbi.1003118.

Li, Heng, Bob Handsaker, Alec Wysoker, Tim Fennell, Jue Ruan, Nils Homer, Gabor Marth, Goncalo Abecasis, Richard Durbin, and 1000 Genome Project Data Processing Subgroup. 2009. “The Sequence Alignment/Map Format and SAMtools.- Bioinformatics (Oxford, England) 25 (16): 2078–79. doi:10.1093/bioinformatics/btp352.

Luo, G., and L. P. Samaranayake. 2002. “Candida Glabrata, an Emerging Fungal Pathogen, Exhibits Superior Relative Cell Surface Hydrophobicity and Adhesion to Denture Acrylic Surfaces Compared with Candida Albicans.- ApMIS: Acta Pathologica, Microbiologica, et Immunologica Scandinavica 110 (9): 601–10.

Maerkl, Sebastian J., and Stephen R. Quake. 2009. “Experimental Determination of the Evolvability of a Transcription Factor.- Proceedings of the National Academy of Sciences of the United States of America 106 (44): 18650–55. doi:10.1073/pnas.0907688106.

McKeown, Alesia N., Jamie T. Bridgham, Dave W. Anderson, Michael N. Murphy, Eric A. Ortlund, and Joseph W. Thornton. 2014. “Evolution of DNA Specificity in a Transcription Factor Family Produced a New Gene Regulatory Module.- Cell 159 (1): 58–68. doi:10.1016/j.cell.2014.09.003.

Murtagh, Fionn, and Pierre Legendre. 2014. “Ward’ s Hierarchical Agglomerative Clustering Method: Which Algorithms Implement Ward’ s Criterion?- Journal of Classification 31 (3): 274–95. doi:10.1007/s00357-014-9161-z.

Nitta, Kazuhiro R., Arttu Jolma, Yimeng Yin, Ekaterina Morgunova, Teemu Kivioja, Junaid Akhtar, Korneel Hens, et al. 2015. “Conservation of Transcription Factor Binding Specificities across 600 Million Years of Bilateria Evolution.- eLife 4 (March): e04837. doi:10.7554/eLife.04837.

Ogawa, Nobuo, Joseph DeRisi, and Patrick O. Brown. 2000. “New Components of a System for Phosphate Accumulation and Polyphosphate Metabolism in Saccharomyces cerevisiaeRevealed by Genomic Expression Analysis.- Molecular Biology of the Cell 11 (12): 4309–21. doi:10.1091/mbc.11.12.4309.

O’Neill, E. M., A. Kaffman, E. R. Jolly, and E. K. O’Shea. 1996. “Regulation of PHO4 Nuclear Localization by the PHO80-PHO85 Cyclin-CDK Complex.- Science (New York, N.Y.) 271 (5246): 209–12.

Orkwis, Brianne R., Danielle L. Davies, Christine L. Kerwin, Dominique Sanglard, and Dennis D. Wykoff. 2010. “Novel Acid Phosphatase in Candida Glabrata Suggests Selective Pressure and Niche Specialization in the Phosphate Signal Transduction Pathway.- Genetics 186 (3): 885–95. doi:10.1534/genetics.110.120824.

Orlando, Kelly A., Christine L. Iosue, Sarah G. Leone, Danielle L. Davies, and Dennis D. Wykoff. 2015. “A Paralogue of the Phosphomutase-like Gene Family in Candida Glabrata, CgPmu2, Gained Broad-Range Phosphatase Activity due to a Small Number of Clustered Substitutions.- The Biochemical Journal 471 (2): 187–98. doi:10.1042/BJ20150611.

Pieterse, Bart, Renger H. Jellema, and Mariët J. van der Werf. 2006. “Quenching of Microbial Samples for Increased Reliability of Microarray Data.- Journal of Microbiological Methods 64 (2): 207–16. doi:10.1016/j.mimet.2005.04.035.

Quinlan, Aaron R., and Ira M. Hall. 2010. “BEDTools: A Flexible Suite of Utilities for Comparing Genomic Features.- Bioinformatics 26 (6): 841–42. doi:10.1093/bioinformatics/btq033.

Reid, Robert J D, Michael Lisby, and Rodney Rothstein. 2002. “Cloning-Free Genome Alterations in Saccharomyces Cerevisiae Using Adaptamer-Mediated PCR.- Methods in Enzymology 350: 258–77.

Rhee, Ho Sung, and B. Franklin Pugh. 2012. “ChIP-Exo Method for Identifying Genomic Location of DNA-Binding Proteins with near-Single-Nucleotide Accuracy.- Current Protocols in Molecular Biology / Edited by Frederick M. Ausubel…[et Al.] Chapter 21 (October): Unit 21.24. doi:10.1002/0471142727.mb2124s100.

Ritchie, Matthew E., Belinda Phipson, Di Wu, Yifang Hu, Charity W. Law, Wei Shi, and Gordon K. Smyth. 2015. “Limma Powers Differential Expression Analyses for RNA-Sequencing and Microarray Studies.- Nucleic Acids Research, January, gkv007. doi:10.1093/nar/gkv007.

Robinson, Mark D., Davis J. McCarthy, and Gordon K. Smyth. 2010. “edgeR: A Bioconductor Package for Differential Expression Analysis of Digital Gene Expression Data.- Bioinformatics 26 (1): 139–40. doi:10.1093/bioinformatics/btp616.

Robinson, Mark D., and Alicia Oshlack. 2010. “A Scaling Normalization Method for Differential Expression Analysis of RNA-Seq Data.- Genome Biology 11: R25. doi:10.1186/gb-2010-11-3-r25.

Romanowski, Kathleen, Alexander Zaborin, Vesta Valuckaite, Ronda J. Rolfes, Trissa Babrowski, Cindy Bethel, Andrea Olivas, Olga Zaborina, and John C. Alverdy. 2012. “Candida Albicans Isolates from the Gut of Critically Ill Patients Respond to Phosphate Limitation by Expressing Filaments and a Lethal Phenotype.- Edited by Neeraj Chauhan. PLoS ONE 7 (1): e30119. doi:10.1371/journal.pone.0030119.

Sayou, Camille, Marie Monniaux, Max H. Nanao, Edwige Moyroud, Samuel F. Brockington, Emmanuel Thévenon, Hicham Chahtane, et al. 2014. “A Promiscuous Intermediate Underlies the Evolution of LEAFY DNA Binding Specificity.- Science 343 (6171): 645–48. doi:10.1126/science.1248229.

Serandour, Aurelien A., Gordon D. Brown, Joshua D. Cohen, and Jason S. Carroll. 2013. “Development of an Illumina-Based ChIP-Exonuclease Method Provides Insight into FoxA1-DnA Binding Properties.- Genome Biology 14 (12): R147. doi:10.1186/gb-2013-14-12-r147.

Shao, D., C. L. Creasy, and L. W. Bergman. 1996. “Interaction of Saccharomyces Cerevisiae Pho2 with Pho4 Increases the Accessibility of the Activation Domain of Pho4.- Molecular and General Genetics MGG 251 (3): 358–64. doi:10.1007/BF02172527.

Slattery, Matthew, Todd Riley, Peng Liu, Namiko Abe, Pilar Gomez-Alcala, Iris Dror, Tianyin Zhou, et al. 2011. “Cofactor Binding Evokes Latent Differences in DNA Binding Specificity between Hox Proteins.- Cell 147 (6): 1270–82. doi:10.1016/j.cell.2011.10.053.

Smyth, Gordon K., Joëlle Michaud, and Hamish S. Scott. 2005. “Use of within-Array Replicate Spots for Assessing Differential Expression in Microarray Experiments.- Bioinformatics (Oxford, England) 21 (9): 2067–75. doi:10.1093/bioinformatics/bti270.

Stern, David L. 2010. Evolution, Development, and the Predictable Genome. 1 edition. Greenwood Village, Colo: W. H. Freeman.

Struhl, Kevin. 1987. “The DNA-Binding Domains of the Jun Oncoprotein and the Yeast GCN4 Transcriptional Activator Protein Are Functionally Homologous.- Cell 50 (6): 841–46. doi:10.1016/0092-8674(87)90511-3.

Thomas-Chollier, Morgane, Carl Herrmann, Matthieu Defrance, Olivier Sand, Denis Thieffry, and Jacques van Helden. 2011. “RSAT Peak-Motifs: Motif Analysis in Full-Size ChIP-Seq Datasets.- Nucleic Acids Research, December, gkr1104. doi:10.1093/nar/gkr1104.

Thompson, Dawn A., Sushmita Roy, Michelle Chan, Mark P. Styczynsky, Jenna Pfiffner, Courtney French, Amanda Socha, et al. 2013. “Evolutionary Principles of Modular Gene Regulation in Yeasts.- eLife 2 (June): e00603. doi:10.7554/eLife.00603.

Vogel, K, W Hörz, and A Hinnen. 1989. “The Two Positively Acting Regulatory Proteins PHO2 and PHO4 Physically Interact with PHO5 Upstream Activation Regions.- Molecular and Cellular Biology 9 (5): 2050–57.

Wapinski, Ilan, Avi Pfeffer, Nir Friedman, and Aviv Regev. 2007. “Natural History and Evolutionary Principles of Gene Duplication in Fungi.- Nature 449 (7158): 54–61. doi:10.1038/nature06107.

Ward, Joe H. Jr. 1963. “Hierarchical Grouping to Optimize an Objective Function.- Journal of the American Statistical Association 58 (301): 236–44. doi:10.1080/01621459.1963.10500845.

Werven, Folkert J. van, and H. Th Marc Timmers. 2006. “The Use of Biotin Tagging in Saccharomyces Cerevisiae Improves the Sensitivity of Chromatin Immunoprecipitation.- Nucleic Acids Research 34 (4): e33–e33. doi:10.1093/nar/gkl003.

Wilson, Michael D., Nuno L. Barbosa-Morais, Dominic Schmidt, Caitlin M. Conboy, Lesley Vanes, Victor L. J. Tybulewicz, Elizabeth M. C. Fisher, Simon Tavaré, and Duncan T. Odom. 2008. “Species-Specific Transcription in Mice Carrying Human Chromosome 21.- Science (New York, N.Y.) 322 (5900): 434–38. doi:10.1126/science.1160930.

Wittkopp, Patricia J., and Gizem Kalay. 2011. “Cis-Regulatory Elements: Molecular Mechanisms and Evolutionary Processes Underlying Divergence.- Nature Reviews. Genetics 13 (1): 59–69. doi:10.1038/nrg3095.

Wykoff, Dennis D., Abbas H. Rizvi, Jonathan M. Raser, Brian Margolin, and Erin K. O’Shea. 2007. “Positive Feedback Regulates Switching of Phosphate Transporters in S. Cerevisiae.- Molecular Cell 27 (6): 1005–13. doi:10.1016/j.molcel.2007.07.022.

Zhang, Yong, Tao Liu, Clifford A. Meyer, Jérôme Eeckhoute, David S. Johnson, Bradley E. Bernstein, Chad Nusbaum, et al. 2008. “Model-Based Analysis of ChIP-Seq (MACS).- Genome Biology 9 (9): R137. doi:10.1186/gb-2008-9-9-r137.

Zhao, Yue, David Granas, and Gary D. Stormo. 2009. “Inferring Binding Energies from Selected Binding Sites.- PLoS Computational Biology 5 (12): e1000590. doi:10.1371/journal.pcbi.1000590.

Zhou, Xu, and Erin K. O’Shea. 2011. “Integrated Approaches Reveal Determinants of Genome-Wide Binding and Function of the Transcription Factor Pho4.- Molecular Cell 42 (6): 826–36. doi:10.1016/j.molcel.2011.05.025.

Zhu, Cong, Kelsey J. R. P. Byers, Rachel Patton McCord, Zhenwei Shi, Michael F. Berger, Daniel E. Newburger, Katrina Saulrieta, et al. 2009. “High-Resolution DNA-Binding Specificity Analysis of Yeast Transcription Factors.- Genome Research 19 (4): 556–66. doi:10.1101/gr.090233.108.

